# A nuclear RNA degradation code for eukaryotic transcriptome surveillance

**DOI:** 10.1101/2024.07.23.604837

**Authors:** Lindsey V. Soles, Liang Liu, Xudong Zou, Yoseop Yoon, Shuangyu Li, Lusong Tian, Marielle Cárdenas Valdez, Angela Yu, Hong Yin, Wei Li, Fangyuan Ding, Georg Seelig, Lei Li, Yongsheng Shi

## Abstract

The RNA exosome plays critical roles in eukaryotic RNA degradation, but it remains unclear how the exosome specifically recognizes its targets. The PAXT connection is an adaptor that recruits the exosome to polyadenylated RNAs in the nucleus, especially transcripts polyadenylated at intronic poly(A) sites. Here we show that PAXT-mediated RNA degradation is induced by the combination of a 5′ splice site and a poly(A) junction, but not by either sequence alone. These sequences are bound by U1 snRNP and cleavage/polyadenylation factors, which in turn cooperatively recruit PAXT. As the 5′ splice site-poly(A) junction combination is typically not found on correctly processed full-length RNAs, we propose that it functions as a “nuclear RNA degradation code” (NRDC). Importantly, disease-associated single nucleotide polymorphisms that create novel 5′ splice sites in 3′ untranslated regions can induce aberrant mRNA degradation via the NRDC mechanism. Together our study identified the first NRDC, revealed its recognition mechanism, and characterized its role in human diseases.

## INTRODUCTION

Eukaryotic genomes are pervasively transcribed^1,2^ and a large portion of these transcripts are quickly degraded in the nucleus, including many long noncoding RNAs (lncRNAs), enhancer RNAs, transcripts of repetitive elements, promoter upstream transcripts (PROMPTs), and misprocessed mRNAs.^3^ Several ribonucleases have been identified that play essential roles in transcriptome surveillance.^4^ Mutations in these proteins cause numerous diseases, especially neurological disorders.^3,5^ The RNA exosome is an essential RNA degradation machine that employs 3′—5′ exoribonucleolytic and endoribonucleolytic enzymes to process and degrade RNAs.^5^ The RNA exosome exhibits weak and non-specific degradation activity on its own and requires adaptor complexes to recognize and efficiently degrade its target substrates.^3,6,7^ Two exosome adaptor complexes have been identified in the nucleoplasm of mammalian cells: the Nuclear Exosome Targeting (NEXT) complex^8^ and the Poly(A) Tail Exosome Targeting (PAXT) connection.^9,10^ NEXT primarily targets non-polyadenylated RNAs and PAXT generally targets polyadenylated RNAs.^11^ However, it remains unclear how these adaptors specifically recognize their respective targets. The core of the PAXT connection is a stable dimer formed by the helicase MTR4, which is also a component of the NEXT complex, and the zinc-finger protein ZFC3H1.^9,10^ The nuclear poly(A) binding protein PABPN1 associates with the PAXT core in a partially RNA-dependent manner and has been suggested to direct PAXT to polyadenylated RNAs.^9^ However, as the vast majority of mRNAs and many lncRNAs have poly(A) tails, additional RNA features must be necessary to specifically recruit the PAXT connection and the RNA exosome for degradation.

The 3′ ends of nearly all eukaryotic mRNAs are formed by cleavage and polyadenylation (CPA).^12,13^ Mammalian poly(A) sites (PASs) typically consist of an A(A/U)UAAA hexamer, called the poly(A) signal, a downstream U/GU-rich element, and other auxiliary sequences. These sequences are recognized by several CPA factors, including CPSF and CstF, to assemble the CPA complex in which the two chemical reactions take place.^14,15^ The majority of human genes produce multiple transcript isoforms by using alternative PASs through a mechanism called alternative polyadenylation.^16,17^ Approximately 30-40% of annotated PASs are within introns and transcripts produced by intronic polyadenylation (IPA) often encode truncated and non-functional proteins.^18–21^ Elevated levels of IPA transcripts are detected in leukemia cells and have been proposed to inactivate tumor suppressor genes.^20^ Recently, an IPA transcript of the *TP53* gene was shown to be oncogenic.^21^ As such, the accumulation of IPA transcripts is generally deleterious and thus must be repressed. Previous studies suggested that the splicing factor U1 snRNP inhibits CPA at intronic PASs through a splicing-independent mechanism called telescripting.^22,23^ More recent studies have provided evidence that at least some IPA transcripts are generated but are targeted for exosome-mediated degradation by PAXT.^10,24^ However, similar to other unstable transcripts, it remains unknown which and how IPA transcripts are specifically recognized for degradation.

In this study, we define the mechanism by which the PAXT connection recognizes specific RNA targets. Our results revealed that, among polyadenylated transcripts of human genes, IPA transcripts are the major targets of PAXT. Importantly, we showed that the combination of a 5′ splice site (ss) and a poly(A) junction, but neither sequence alone, triggers RNA degradation. These sequences are recognized by U1 snRNP and CPA factors, which in turn synergistically recruit PAXT and the exosome. As 5′ splice sites and poly(A) junctions are typically found in unspliced pre-mRNAs and processed mRNAs respectively, we propose that the abnormal presence of both sequences on IPA transcripts constitutes a “nuclear RNA degradation code” (NRDC) that marks them for degradation. Finally, we demonstrate that many disease-associated single nucleotide polymorphisms (SNPs) create a novel 5′ ss in the 3′ untranslated region, thereby promoting RNA degradation through the NRDC mechanism.

## RESULTS

### IPA transcripts are major targets of PAXT- and exosome-mediated degradation

To comprehensively identify the polyadenylated transcripts that are targeted by the RNA exosome and the PAXT connection, we individually depleted the nuclear exosome factor EXOSC3/RRP40, or the PAXT components MTR4, ZFC3H1, and PABPN1 by RNAi and performed PAS-seq analyses (Fig. S1A).^25,26^ PAS-seq is a 3′-end RNA-sequencing method that not only maps PASs (Fig. S1B), but also measures the relative levels of polyadenylated RNAs. As previous studies have characterized PAXT-targeted PROMPTs in great detail,^9,11^ we have focused on the sense-strand polyadenylated transcripts of all annotated genes. Our results revealed that depletion of EXOSC3, MTR4, or ZFC3H1 led to significantly higher levels of 2,371, 2,874, and 2,076 transcripts respectively (log_2_FC (fold change) > 1, FDR < 0.05). We categorized these transcripts into multiple groups, including those polyadenylated in last exons, upstream internal exons, alternative last exons, or introns. Among the transcripts significantly upregulated in EXOSC3-, MTR4-, and ZFC3H1-depleted cells, IPA transcripts accounted for 43%, 50.7%, and 55.8% respectively (Fig. S1B-C), representing the largest group in all samples. In comparison, PABPN1 depletion caused significantly accumulation of 1,810 transcripts and only 31.5% of them were IPA transcripts (Fig. S1C). More transcripts polyadenylated in the last exons were significantly upregulated upon PABPN1 knockdown (51.4%, Fig. S1C), suggesting that the effect of PABPN1 depletion was distinct from that of EXOSC3, MTR4, or ZFC3H1 depletion. As IPA transcripts seemed to be the major group regulated by the exosome and ZFC3H1/MTR4, we compared the change in RNA levels between IPA and non-IPA transcripts, including those polyadenylated in last exons, upstream internal exons or alternative last exons (Fig. S1B). As shown in Fig. 1A, a much greater proportion of IPA transcripts accumulated to high levels in EXOSC3-, MTR4- and ZFC3H1-depleted cells compared to non-IPA transcripts. In contrast to these largely unidirectional changes, depletion of PABPN1 caused significant changes in IPA transcript abundance in both directions, (Fig. 1A). Two example genes are shown to illustrate these changes (Fig. 1B-C). For the tumor suppressor gene *TP53* and the gene *RNF166*, the latter of which is implicated in cancer and immunity,^27–29^ very low levels of IPA transcripts were detected in control cells while depletion of EXOSC3, MTR4, and ZFC3H1 led to a significant increase in the abundance of their IPA transcripts. While PABPN1 depletion also resulted in significant increases in their IPA transcript abundance, the magnitude was more modest (Fig. 1B-C). Hierarchical clustering of all IPA transcripts that were significantly upregulated in at least one condition revealed that ZFC3H1- and MTR4-depletion samples clustered together (Fig. 1D). This cluster was closely related to EXOSC3-depletion but was more distinct from the PABPN1-depletion sample (Fig. 1D). These results are consistent with reports that ZFC3H1 and MTR4 function as a tight core complex in the PAXT connection while the association between PABPN1 and the ZFC3H1/MTR4 core is more dynamic and partially RNA-dependent.^5^ Together, our PAS-seq results suggest that, among all polyadenylated transcripts of annotated human genes, IPA transcripts are the major targets of PAXT-mediated degradation by the RNA exosome and that the ZFC3H1/MTR4 dimer plays a prominent role in their regulation.

**Figure 1.**
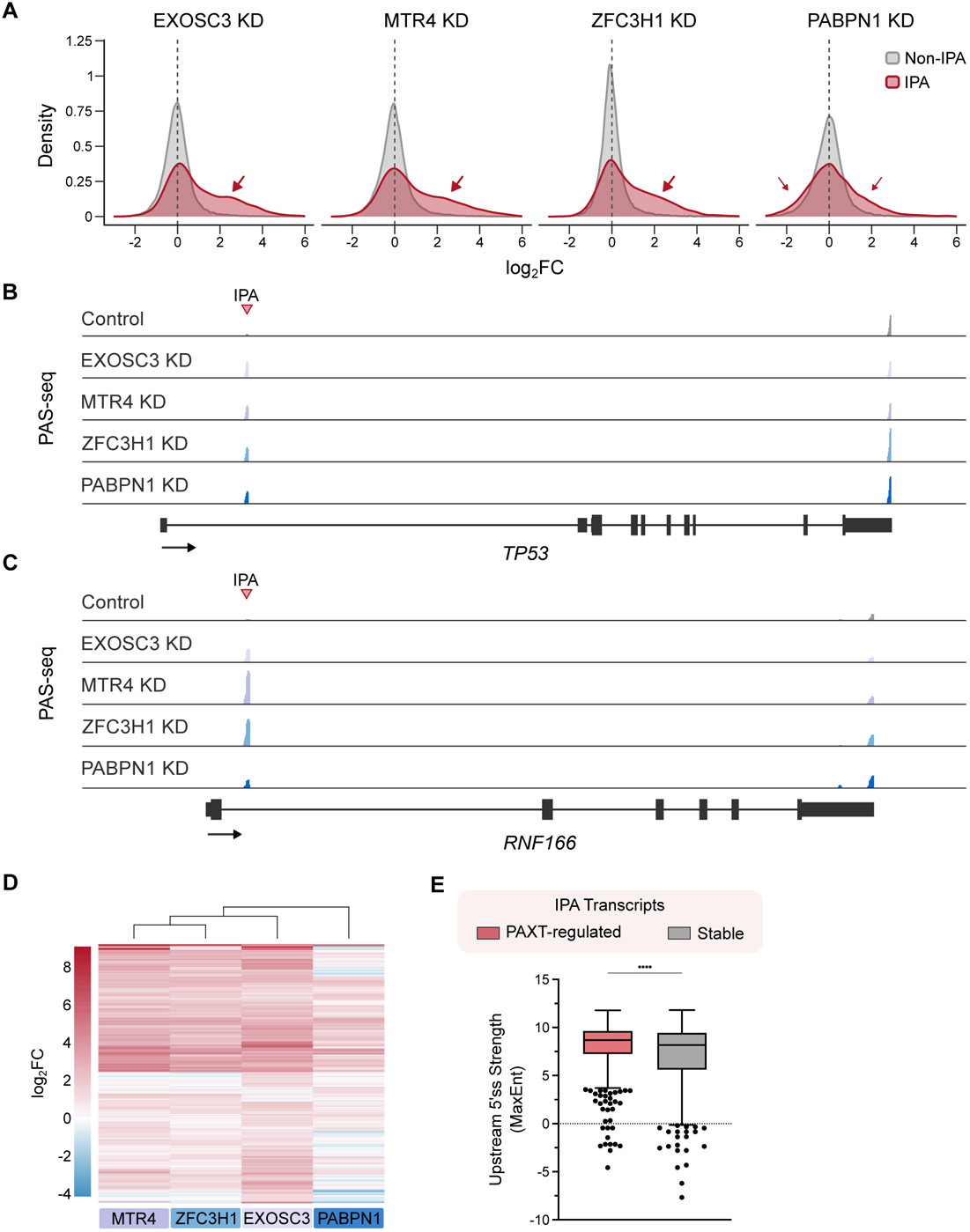
IPA transcripts are major targets of PAXT-mediated degradation. **A**, Density plots of the log_2_FC for all IPA or non-IPA transcripts following depletion of EXOSC3, MTR4, ZFC3H1, or PABPN1 in wild-type HEK293T cells. IPA transcripts are depicted in red and non-IPA transcripts are depicted in grey. A positive log_2_FC indicates that the transcript is upregulated upon knockdown. **B** & **C**, PAS-seq data tracks for the genes *TP53* (**B**) and *RNF166* (**C**) following depletion of EXOSC3, MTR4, ZFC3H1, or PABPN1 by RNAi in wild-type HEK293T cells. Each peak represents reads mapped to the 3′ end of mRNAs. **D**, Heatmap of the log_2_FC measured for all IPA transcripts that were found to be upregulated in at least one knockdown condition (log_2_FC > 1, FDR ≤ 0.05). A positive log_2_FC indicates that the IPA transcript is upregulated upon knockdown. **E**, Box plot depicting the strength of the nearest upstream 5′ ss as measured by MaxEnt. PAXT-regulated: Significantly upregulated IPA transcripts (log_2_FC > 1, FDR ≤ 0.05, *N* = 507) following depletion of EXOSC3, MTR4, and ZFC3H1. Stable IPA transcripts: IPA transcripts whose expression did not significantly change following depletion of EXOSC3, MTR4, and ZFC3H1 (Counts per million (CPM) in control cells > 1, log_2_FC < 1 and log_2_FC > -1, FDR > 0.05, *N* = 687). Statistical analysis was calculated by Mann-Whitney test.

While a large portion of IPA transcripts accumulated to significantly higher levels in EXOSC3-, MTR4-, or ZFC3H1-depleted cells (Fig. 1A, marked by arrows), some remain unchanged. To characterize the molecular basis for such differences in IPA transcript stabilities, we compared the 5′ ss strength, as measured by the MaxEnt score,^30^ of PAXT-regulated unstable IPA transcripts (log_2_FC > 1, FDR ≤ 0.05, in EXOSC3-, MTR4-, and ZFC3H1-depleted cells, N = 507) and stable IPA transcripts (Counts per million (CPM) in control cells > 1, -1 < log_2_FC < 1, FDR > 0.05, N = 687). We found that the 5′ ss was significantly stronger for PAXT-regulated IPA transcripts (median: 8.69) than for stable IPA transcripts (median: 8.17) (p < 0.0001, Mann-Whitney test) (Fig. 1E), indicating that stronger U1 snRNP binding to the upstream 5′ ss on IPA transcripts correlates with more efficient PAXT-mediated degradation by the exosome.

### Sequence determinants of exosomal degradation of IPA transcripts

How are IPA transcripts specifically recognized by the exosome? As all IPA transcripts share two common features, a 5′ ss and a downstream poly(A) tail (Fig. S1B), we tested their roles in degradation using a series of reporters. To mimic IPA transcripts, we inserted a 5′ ss sequence from the *NXF1* gene, which was previously shown to efficiently bind U1 snRNP,^31^ into the 3′ untranslated region (UTR) of an eGFP reporter (Fig. 2A). To test the role of the 5′ ss, we used the wild-type (WT) or a non-functional mutant (Mut) form of the 5′ ss. To test the impact of the poly(A) tail, we included in these reporters either a PAS (the PAS of the bovine growth hormone (*bGH*) gene or the adenovirus major late transcript (L3)), or a sequence that conferred a non-polyadenylated 3′ end (the 3′ end sequence of the nuclear lncRNA *MALAT1* or the 3′ end sequence from the replication-dependent histone gene *H2AC18*) (Fig. 2A). The 3′ end of *MALAT1* RNA is formed by RNase P-mediated cleavage and forms a stable RNA triple helix.^32^ In contrast, the 3′ end of *H2AC18* mRNA is formed via a single endoribonucleolytic cleavage reaction downstream of a RNA stem-loop structure.^33^ As a result, the 3′ ends of both *MALAT1* and *H2AC18* lack a poly(A) tail. These reporters were transfected into HEK293T cells and their mRNA levels relative to those of a co-transfected control reporter were measured by RT-qPCR. Strikingly, we observed that the presence of a 5′ ss in the 3′ UTR of a polyadenylated reporter caused significant decreases in the mRNA levels (4.0-fold for the *bGH* PAS-containing reporter, p < 0.0001, unpaired t-test; 7.6-fold for the L3 PAS-containing reporter, p < 0.0001, unpaired t-test) (Fig. 2B). The 5′ ss induced only a mild decrease (1.6-fold, p = 0.0125, unpaired t-test) in the reporter with the histone *H2AC18* 3′ end, consistent with a previous study.^34^ In contrast, a 5′ ss induced a mild increase in the mRNA levels in the reporter with the *MALAT1* 3′ end (1.5-fold, p = 0.0007, unpaired t-test, Fig. 2B). These results suggest that both a 5′ ss and a poly(A) tail are required to efficiently repress reporter mRNA levels.

**Figure 2.**
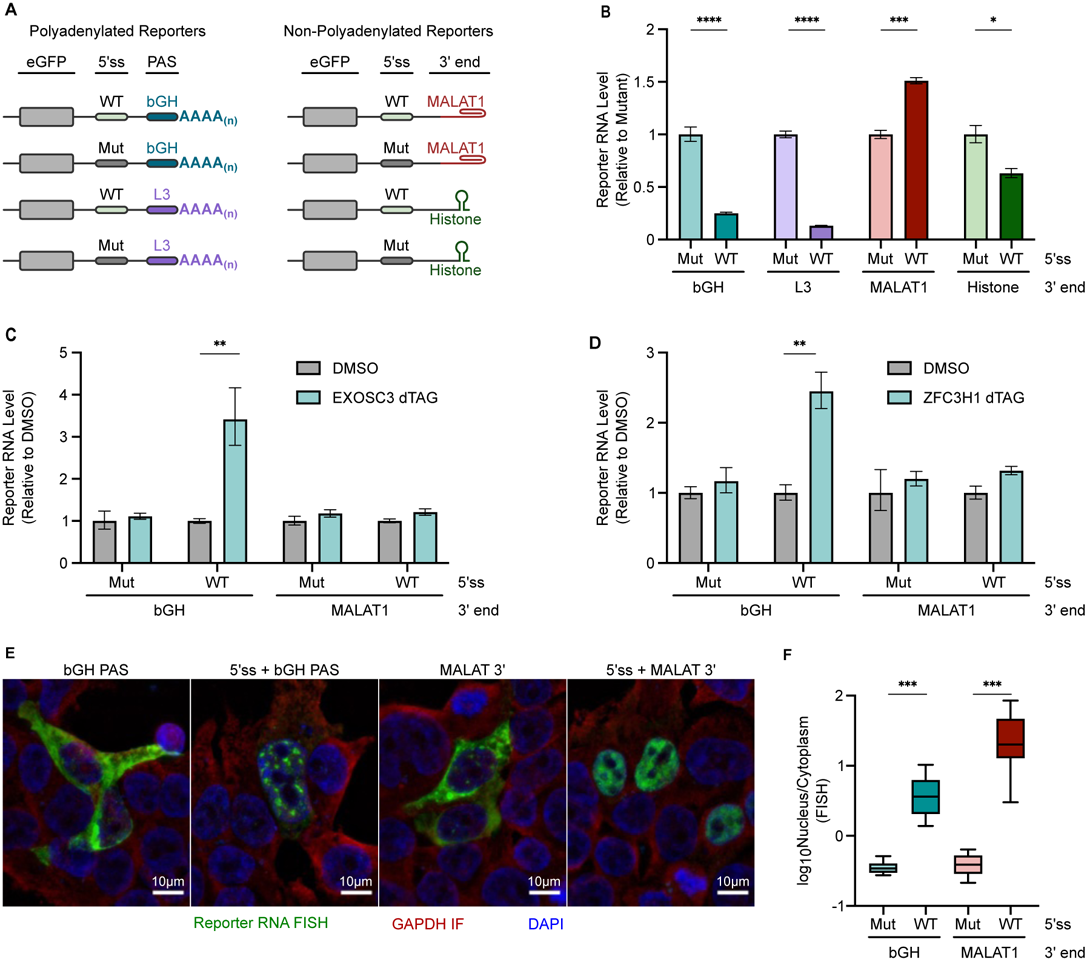
A 5′ splice site and a poly(A) tail trigger PAXT-dependent exosomal degradation. **A**, Schematic of the mRNA structure of the eGFP reporters used in **B** - **F**. The reporters contain the coding sequence for eGFP followed by either a wild-type (WT) or mutant (Mut) 5′ ss in the 3′ UTR. The 3′ end of the reporter includes a poly(A) site (PAS), the 3′ end sequence from *MALAT1* (MALAT1), or the 3′ end sequence from *H2AC18* (Histone) (see Methods for details). **B**, RT-qPCR analysis of random-primed cDNA following overexpression of each reporter. The reporter mRNA levels were normalized to the expression of a co-transfected renilla luciferase control plasmid. Then for each 3′ end pair, the mRNA level of the 5′ ss wild-type reporter was calculated relative to the 5′ ss mutant reporter. Statistical analysis from *n* = 3 independent samples was calculated using unpaired t-tests. **C** & **D**, RT-qPCR analysis of random-primed cDNA following overexpression of each reporter and dTAG-induced depletion of EXOSC3 (**C**) or ZFC3H1 (**D**). The reporter mRNA levels were normalized to the expression of a co-transfected firefly luciferase control plasmid. Then for each reporter, the mRNA level in the dTAG condition was calculated relative to the DMSO control condition. Statistical analysis from *n* = 3 independent samples was calculated using unpaired t-tests. **E**, Combined reporter RNA FISH and GAPDH immunofluorescence (IF) in wild-type HEK293T cells. The reporter RNA FISH signal is shown in green. GAPDH IF signal is shown in red. Nuclei were stained using DAPI and are depicted in blue. **F**, Boxplot displaying the quantification of the reporter RNA FISH signal in the nucleus versus cytoplasm in **E** (see Methods). For each 3′ end, the Nucleus/Cytoplasm FISH signal ratio of the 5′ wild-type reporter was normalized to the Nucleus/Cytoplasm FISH signal ratio of the 5′ ss mutant reporter. Y-axis values are formatted as log values. Statistical analysis from *n* = 10 (5′ ss-*bGH*, 5′ss Mut-*bGH*), *n* = 12 (5′ss-*MALAT1* 3′ end), or *n* = 11 (5′ss Mut-*MALAT1* 3′ end) images per reporter was calculated using unpaired t-tests.

The 5′ ss and PAS could repress reporter mRNA levels by inhibiting RNA synthesis, for example by blocking CPA via telescripting^22^ and/or by promoting RNA degradation. To distinguish between these possibilities, we performed reporter assays in control or exosome-depleted cells. To deplete the exosome, we generated a HEK293T cell line in which an inducible degron tag (dTAG)^35^ was fused to the C-terminus of the exosome subunit EXOSC3 via CRISPR/Cas9-mediated genome editing. Treatment of this cell line with the dTAG molecule for 8 hours led to near complete degradation of the EXOSC3 protein by the ubiquitin-proteosome system (Fig. S2A). We transfected reporters that contained a WT or a Mut 5′ ss and the *bGH* PAS or the *MALAT1* 3′ end sequence into the mock (DMSO) or dTAG-treated EXOSC3 degron cell line and compared the reporter mRNA levels. Compared to mock-treated cells, EXOSC3 depletion led to a significant increase in the mRNA level of the 5′ ss-*bGH* PAS reporter (3.4-fold, p = 0.004, unpaired t-test), but not for other reporters (Fig. 2C). In fact, in EXOSC3-depleted cells, the mRNA levels of the 5′ ss-*bGH* PAS reporter accumulated to a similar level to that of the reporter that contained a Mut 5′ ss (compare Fig. 2C and Fig. 2B). These results suggest that a 5′ ss and a poly(A) tail repressed the reporter mRNA levels primarily by promoting exosome-mediated RNA degradation. Next, we sought to determine if such degradation was mediated by the PAXT connection. To this end, we fused a dTAG to the N-terminus of ZFC3H1 (Fig. S2B). dTAG-induced depletion of ZFC3H1 caused a significant increase in the mRNA levels of the 5′ ss-PAS reporter (2.4-fold, p = 0.0041, unpaired t-test), but not for other reporters (Fig. 2D), suggesting that the 5′ ss and poly(A) tail-induced exosomal degradation was indeed mediated by PAXT. Note that the increase in mRNA levels in ZFC3H1-depleted cells was lower than that observed in EXOSC3-depleted cells (2.4-fold vs. 3.4-fold, Fig. 2C-D). One likely reason for this discrepancy is that ZFC3H1 has multiple isoforms and the dTAG treatment primarily depleted the largest isoform (Fig. S2B).

The essential role of a 5′ ss in inducing RNA degradation suggests that U1 snRNP is required for this process. To test if U1 snRNP-binding is sufficient, we artificially recruited U1 snRNP to a 3′ UTR sequence that did not contain a 5′ ss and monitored its effect on RNA stability. To this end, we overexpressed a Mut U1 snRNA that can hybridize to the Mut 5′ ss in the *bGH* PAS-containing reporter (Fig. S2C).^36^ Compared to the control non-targeting U1 snRNA, overexpression of the Mut U1 snRNA led to a 1.8-fold decrease in the Mut 5′ ss-containing mRNA level (p = 0.0023, unpaired t-test) (Fig. S2D), suggesting that targeting U1 snRNP to the 3′ UTR is sufficient to induce RNA degradation.

We next investigated how a 5′ ss and a poly(A) tail impacted the subcellular localization of the reporter mRNAs by performing RNA Fluorescence In Situ Hybridization (RNA-FISH). Without a 5′ ss in their 3′ UTR, the reporter mRNAs that contained either a poly(A) tail or the *MALAT1* 3′ end primarily localized to the cytoplasm (Fig. 2E-F), suggesting that the mRNAs from both reporters were efficiently exported. In contrast, the mRNAs of both 5′ ss-containing reporters were retained in the nucleus (Fig. 2E-F). This is consistent with previous studies showing that a 5′ ss promotes nuclear retention of both mRNAs and non-coding RNAs.^24,31,37^ In line with the nuclear sequestration of their mRNAs, the eGFP protein levels of all 5′ ss-containing reporters were significantly lower compared to their counterparts without a 5′ ss (Fig. S2F-H). Interestingly, although the mRNAs of both 5′ ss-PAS and 5′ ss-*MALAT1* 3′ end reporters were retained in the nucleus, their distribution patterns differed. The mRNAs of the 5′ ss-*MALAT1* 3′ end reporter were uniformly distributed in the nucleoplasm (Fig. 2E, fourth panel). In contrast, the 5′ ss-PAS reporter mRNAs were highly concentrated in a number of large nuclear puncta (Fig. 2E, second panel), which partially overlapped with nuclear speckles (Fig. S2E). Together these results suggest that, while a 5′ ss alone induces RNA nuclear retention, the combination of a 5′ ss and a poly(A) tail promotes nuclear retention and PAXT-mediated exosomal degradation.

### U1 snRNP and CPA factors cooperatively recruit PAXT and the exosome

We considered two potential models that may explain how the combination of a 5′ ss and a poly(A) tail promotes PAXT-dependent exosomal RNA degradation. First, previous studies and our results demonstrated that U1 snRNP binds to the 5′ ss and promotes RNA nuclear retention.^24,31,37^ A “nuclear timer model” posits that the prolonged residence of an RNA molecule in the nucleus allows more time for non-sequence-specific exosomal degradation.^5^ Alternatively, U1 snRNP and PABPN1 bound to the 5′ ss and poly(A) tail could cooperatively recruit the PAXT connection and, in turn, the nuclear RNA exosome, leading to sequence-specific degradation. To investigate these possibilities, we performed *in vitro* RNA pulldown assays (Fig. 3A). RNAs with a WT or Mut 5′ ss sequence upstream of the L3 PAS were synthesized by *in vitro* transcription. Three copies of the bacteriophage MS2 hairpin were fused to the 5′ end of the RNAs to allow for pulldown using a fusion protein comprised of the MS2 coat protein (MCP) and the maltose-binding protein (MBP).^38^ The 3xMS2-fused RNAs were first incubated with the MCP-MBP fusion protein and nuclear extract from HeLa cells to allow for protein-RNA interactions and the CPA reactions to occur. As a negative control, a Mut L3 PAS was used in which the poly(A) signal was mutated from AAUAAA to AAGAAA. As shown in Fig. 3B, CPA occurred for RNAs that contained the WT L3 PAS but not the Mut PAS (Fig. 3B, compare lanes 1-6 with lanes 7-12), and the presence of an upstream 5′ ss did not have a significant effect on the reaction efficiency (Fig. 3B, compare lanes 8 and 11). Interestingly, upon longer incubation (150 minutes), the polyadenylated RNA species shifted higher for the 5′ ss-containing RNA (Fig. 3B, compare lanes 12 and 9), suggesting that the 5′ ss induced hyperadenylation. Although the functional significance of such hyperadenylation is currently unclear, it is consistent with previous reports that PAXT and PABPN1-dependent degradation target RNAs are hyperadenylated.^10,39^

**Figure 3.**
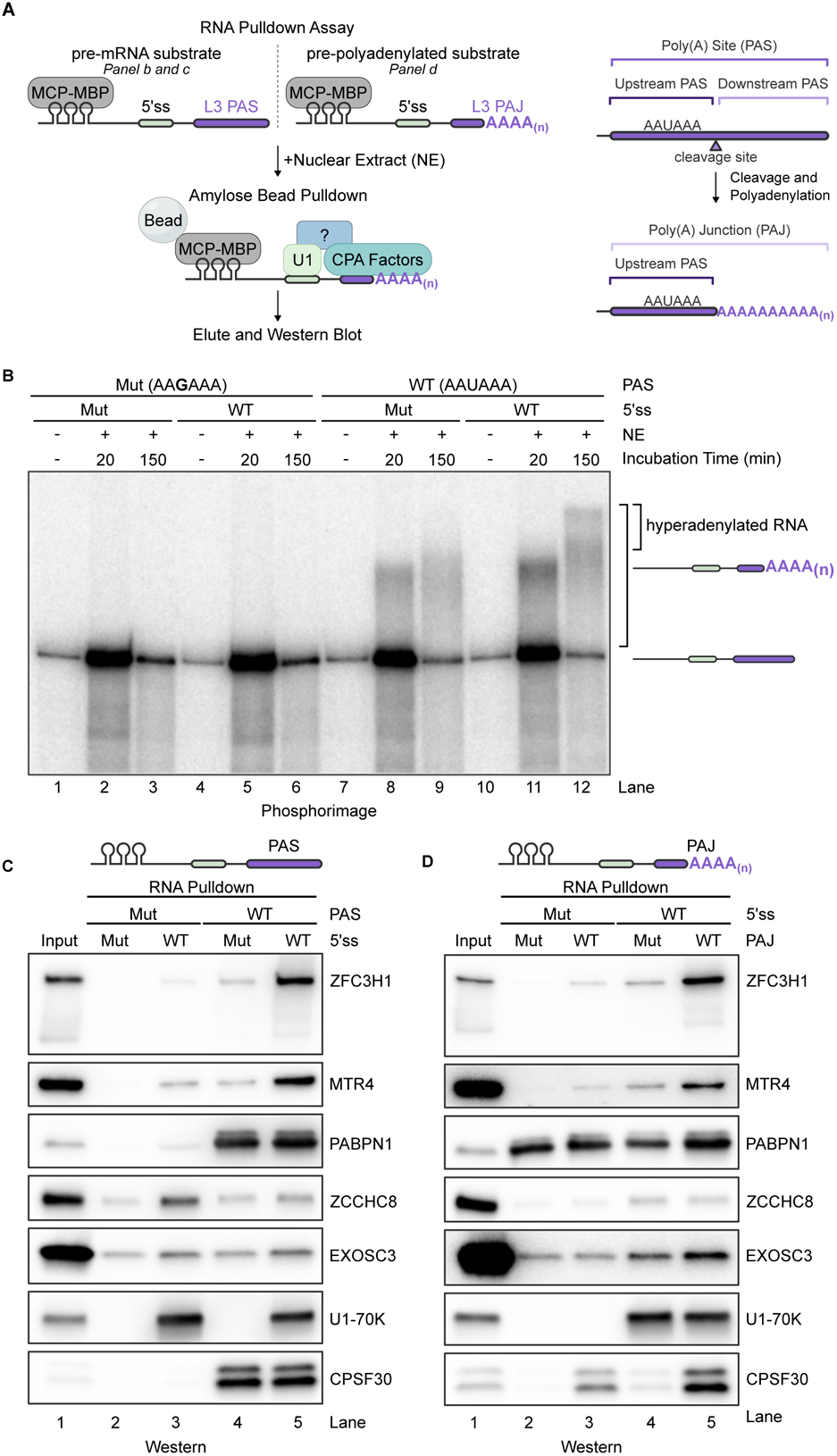
U1 snRNP and cleavage and polyadenylation factors cooperatively recruit PAXT and the RNA exosome. **A**, Left: Schematic of the RNA substrates and the RNA pulldown assays performed. The depicted pre-mRNA substrate is used in **B** & **C** and the pre-polyadenylated substrate is used in **D**. Right: Schematic depicting terms used to describe RNA regions present before and after pre-mRNA cleavage and polyadenylation. **B**, Polyadenylation assay using radiolabeled pre-mRNA substrates. **C**, Western blotting analyses of RNA pulldown assays detecting proteins that assembled on the pre-mRNA substrates. **D**, Western blotting analyses of RNA pulldown assays detecting proteins that assembled on the pre-polyadenylated substrates.

We next pulled down these RNAs using amylose beads and examined the associated proteins by western blotting. As expected, the WT L3 PAS-containing RNAs specifically pulled down CPA factors, including CPSF30 and PABPN1 (Fig. 3C, lanes 4 and 5), and the U1 snRNP component U1-70K specifically associated with WT 5′ ss-containing RNAs (Fig. 3C, lanes 3 and 5), demonstrating the specificity of this assay. Interestingly, both WT 5′ ss-Mut PAS and Mut 5′ ss-WT PAS RNAs were weakly associated with PAXT core subunits, ZFC3H1 and MTR4 (Fig. 3C, lanes 3 and 4). Strikingly, however, dramatically higher amounts of ZFC3H1 and MTR4 were precipitated with the RNAs that contained both a WT 5′ ss and a WT PAS (Fig. 3C, compare lane 5 with lanes 3-4). In contrast, ZCCHC8, a component of the NEXT adaptor complex, was not significantly precipitated (Fig. 3C). These results strongly suggest that both U1 snRNP and CPA factors bind weakly to PAXT individually, but when bound to a 5′ ss and a poly(A) junction on the same RNA molecule, they can synergistically recruit PAXT.

To test if the CPA reaction itself was required for PAXT recruitment, we synthesized pre-polyadenylated versions of the same RNA substrates and performed RNA pulldown assays (Fig. 3A). To do so, we synthesized shorter versions of the PAS-containing RNAs that ended at the natural cleavage sites by *in vitro* transcription and polyadenylated them using *E.coli* poly(A) polymerase. As such, each pre-polyadenylated RNA substrate contained a poly(A) junction (PAJ), which includes the upstream PAS region and the poly(A) tail (Fig. 3A, right panel). To test the role of the poly(A) signal itself, we generated pre-polyadenylated RNAs with a WT (AAUAAA) or Mut (AAGAAA) hexamer. We confirmed that the pre-polyadenylated RNA substrates were of comparable length and that the poly(A) tails were not further extended after incubation in nuclear extract (Fig. S3A-B). These RNAs were incubated with HeLa nuclear extract and subsequently pulled down using amylose beads. As all of the RNAs were pre-polyadenylated, PABPN1 was precipitated with all of them (Fig. 3D). For the RNAs that contained a Mut 5′ ss and a Mut poly(A) signal, neither ZFC3H1 nor MTR4 was precipitated (Fig. 3D, lane 2), demonstrating that PABPN1 alone was not sufficient to recruit PAXT. As expected, the WT 5′ ss-containing RNAs pulled down the U1 snRNP component U1-70K and the WT poly(A) signal-containing RNAs precipitated CPA factors, such as CPSF30 (Fig. 3D). Interestingly, the RNAs containing a Mut 5′ ss-WT poly(A) signal or those containing a WT 5′ ss-Mut poly(A) signal only pulled down low amounts of ZFC3H1 and MTR4 (Fig. 3D, lanes 3 and 4), similar to the corresponding full-length pre-mRNA substrates (Fig. 3C, lanes 3 and 4). In contrast, the polyadenylated RNAs that contained both a WT 5′ ss and a WT poly(A) signal pulled down much higher amounts of ZFC3H1, MTR4, and EXOSC3 (Fig. 3D, compare lane 5 to lanes 3-4). Together both the PAS-containing and the pre-polyadenylated RNA pulldown results demonstrate that both 5′ ss-bound U1 snRNP or poly(A) junction-bound CPA factors displayed weak interactions with PAXT. In contrast, when U1 snRNP and CPA factors bind to their cognate sequences on the same RNA, they synergistically recruit PAXT. Thus, our results identified the 5′ ss-poly(A) junction as the first sequence combination that specifically recruits PAXT for exosomal RNA degradation. Furthermore, our data demonstrated that PABPN1 alone is not sufficient for recruiting PAXT and revealed an essential role for CPA factors, especially CPSF, in this process.

### ZFC3H1 interacts with U1 snRNP and CPA factors

Our RNA pulldown results suggest that both U1 snRNP and CPA factors can weakly associate with PAXT. However, when these factors are bound to the same RNA molecule, they can synergistically recruit PAXT. Consistent with this model, U1-70K and CPA factors, such as subunits of the CPSF complex (CPSF160, CPSF100, and CPSF73), were detected in the interactome of ZFC3H1,^9,24^ but not in that of MTR4.^9^ ZFC3H1 contains four major regions/domains: an acidic short linear motif at the N-terminus called the SLiM region, five coiled coil (CC) domains, a zinc finger domain (ZnF), and tetratricopeptide repeats (TPR) near its C-terminus (Fig. 4A). We next sought to characterize the interactions between ZFC3H1 and U1 snRNP or CPA factors in detail. To this end, we expressed FLAG-tagged ZFC3H1 in HEK293T cells and performed immunoprecipitations using anti-FLAG antibodies (FLAG-IP) (Fig. 4B). Consistent with previous reports, PABPN1, MTR4, and the cap binding complex interacting protein ARS2 were co-immunoprecipitated with ZFC3H1 (Fig. 4B, lanes 6 and 7).^9,40^ ZFC3H1 interacted with MTR4 and ARS2 in an RNase A-resistant manner and, consistent with previous studies,^9^ its association with PABPN1 was severely reduced upon RNase treatment (Fig. 4B, lanes 6 and 7). Low levels of U1 snRNP subunits, including U1-70K, U1A, and U1C as well as CPSF components, including CPSF100, CPSF73, and CPSF30 were co-immunoprecipitated with ZFC3H1 in a partially RNA-dependent manner (Fig. 4B, lanes 6 and 7). These data suggest that ZFC3H1 can indeed interact with both U1 snRNP and CPSF.

**Figure 4.**
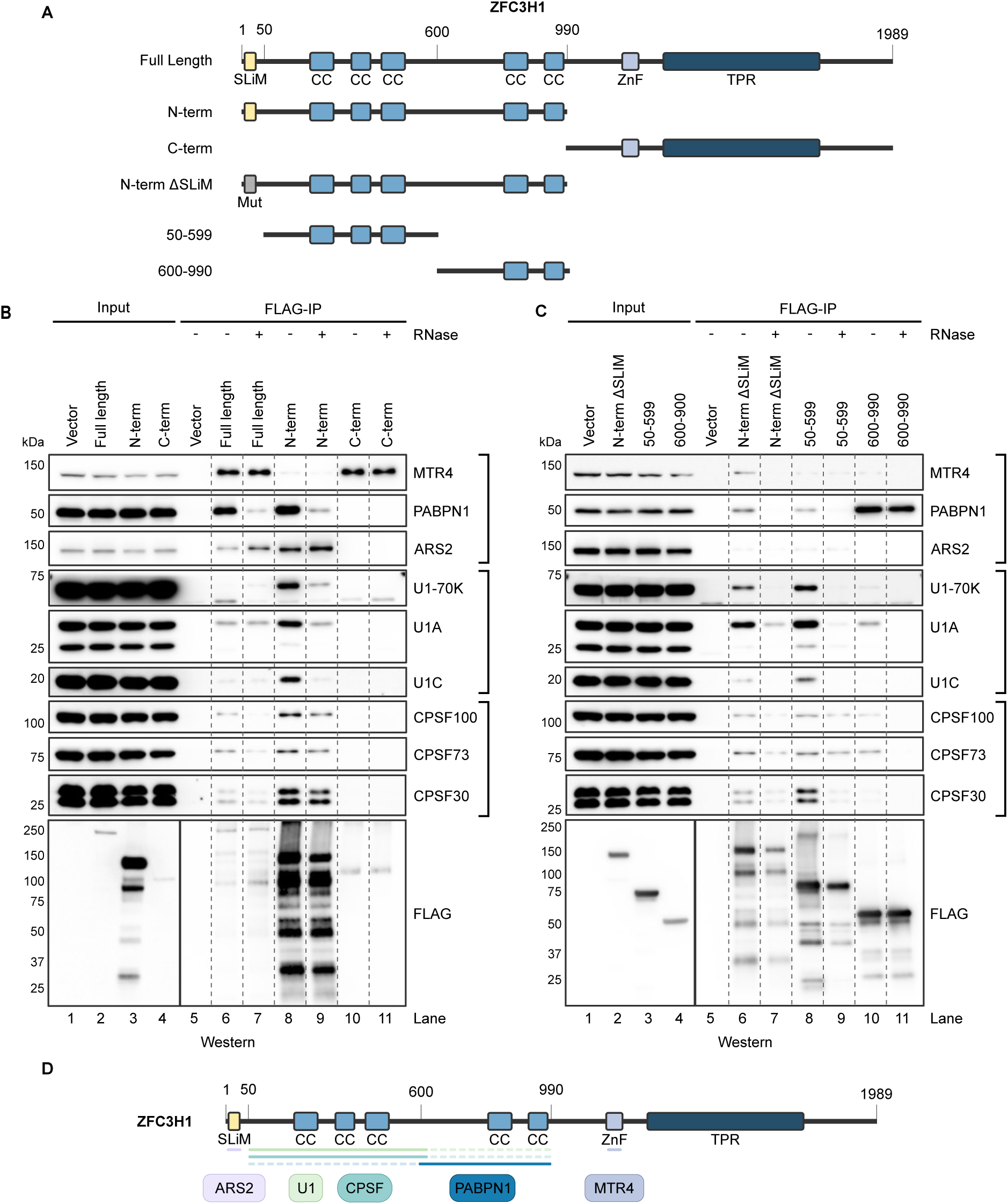
ZFC3H1 interacts with U1 snRNP and cleavage and polyadenylation factors. **A**, Schematic of the predicted domains within full-length ZFC3H1 and the expression constructs used to perform the FLAG-Immunoprecipitation experiments analyzed in **B** & **C**. Relevant residues are indicated as numbers above full-length ZFC3H1. The yellow rectangle represents the short acid-rich linear motif (SLiM) region. The mutated SLiM region within the N-term SLiM mutant (N-term ΔSLiM) construct is represented as a grey rectangle labeled Mut. The light blue rectangles depict coiled-coil (CC) regions. The light purple rectangle represents the zinc-finger domain (ZnF). The dark blue rectangle represents tetratricopeptide repeats (TPR). **B** & **C**, Western blotting analyses of proteins that co-immunoprecipitated with the indicated ZFC3H1 expression construct. The anti-FLAG western blots contain 1% input and 10% eluted IP sample. All other western blots contain 0.5% input and 30% eluted IP sample. **D**, Schematic depicting the interactions between ZFC3H1 and the associated proteins ARS2, U1 snRNP components (U1), subunits of CPSF (CPSF), PABPN1, and MTR4.

To map the regions in ZFC3H1 that mediate interactions with U1 snRNP and CPSF, we performed FLAG-IPs using ZFC3H1 mutants in which a specific region was deleted (Fig. 4A). To broadly define the regions in ZFC3H1 that mediate interactions with U1 snRNP and CPSF, we overexpressed FLAG-tagged N-terminal (residues 1-990) or C-terminal (residues 991-1989) regions of ZFC3H1 and performed IPs with anti-FLAG antibodies. As previously reported,^40^ ARS2 and PABPN1 were co-precipitated with the N-terminal region while MTR4 specifically associated with the C-terminal region of ZFC3H1 (Fig. 4B, lanes 8 and 9, lanes 10 and 11).^40^ Both U1 snRNP and CPSF associated specifically with the N-terminal region of ZFC3H1 (Fig. 4B, lanes 8-11). Notably, the interaction between U1 snRNP and the ZFC3H1 N-terminal region appeared to depend more on RNA than that of CPSF and the ZFC3H1 N-terminal region (Fig. 4B, compare lanes 8 and 9).

We next further dissected the N-terminal region of ZFC3H1 and characterized the role of the SLiM region and the CC domains in mediating interactions with U1 snRNP and CPSF. We first mutated the SLiM region present within residues 12-33 of ZFC3H1 (abbreviated ϕλSLiM, Fig. 4A). The SLiM region contains a highly acidic motif (EEGELEDGEI) and, consistent with previous studies,^41,42^ we observed that substituting four of the acidic residues in the SLiM motif with alanine (EAGALEAGAI) abolished the interaction between ZFC3H1 and ARS2 (Fig. 4C, lanes 6-7, compare with Fig. 4B, lanes 6-7). In contrast, these mutations in the SLiM region did not affect ZFC3H1 interactions with U1 snRNP, CPSF, or PABPN1 (Fig. 4C, lanes 6 and 7), suggesting that the SLiM motif is not necessary for ZFC3H1 to interact with these factors.

We then focused on the N-terminal region of ZFC3H1 that contains five CC domains (residues 50-990). We expressed two FLAG-tagged sub-regions that contained three and two CC domains (residues 50-599 and 600-990, respectively) and performed FLAG IPs (Fig. 4A). Subunits of both U1 snRNP and CPSF components interacted most strongly with residues 50-599 of ZFC3H1 (Fig. 4C, compare lanes 8 and 9 with 10 and 11). The interactions between residues 50-599 of ZFC3H1 and U1 snRNP components were sensitive to RNase A treatment, while the interactions with CPSF components were partially resistant (Fig. 4C, compare lanes 8 and 9). In contrast, PABPN1 interacted most strongly with residues 600-990 of ZFC3H1, consistent with previous reports (Fig. 4C, compare lanes 8 and 9 with lanes 10 and 11).^40^ As summarized in Fig. 4D, our results demonstrated that four distinct regions of ZFC3H1 mediate its interactions with different partners: the SLiM region of ZFC3H1 binds ARS2 (Fig. 4B-C);^41,42^ U1 snRNP and CPSF most strongly interact with residues 50-599 of ZFC3H1 (Fig. 4C); PABPN1 predominantly associates with residues 600-990 of ZFC3H1 (Fig. 4C); MTR4 interacts with the C-terminal zinc finger domain of ZFC3H1 (Fig. 4B).^40^ Importantly, these results demonstrate that both U1 snRNP and CPSF associate with ZFC3H1 and these interactions are mediated by the same region in ZFC3H1 (Fig. 4D). The implications of this seemingly overlapping binding pattern for the cooperative recruitment of ZFC3H1 by U1 snRNP and CPSF are discussed later.

### The nuclear RNA degradation code in human diseases

Our results demonstrate that the combination of a 5′ ss and a poly(A) junction targets RNAs for PAXT-mediated exosomal degradation, and that this combination, which we refer to as the NRDC, plays an important role in specifically degrading IPA transcripts. We next aimed to determine whether and how this mechanism may contribute to human diseases. Previous studies identified a SNP in the 3′ UTR of the human gene *Factor IX* (*F9*), which encodes a blood coagulation factor, that caused significant downregulation of its protein levels and, in turn, severe hemophilia B.^43^ This SNP created a novel 5′ ss by increasing the 5′ ss MaxEnt score of the surrounding sequence from 2.75 in the reference genome sequence to 10.93 in the SNP-containing sequence, which is higher than the median 5′ ss strength across all human introns (mean: 8.66) (Fig. 5A). This novel SNP-created 5′ ss in the 3′ UTR is located 208 nucleotides upstream of the poly(A) signal and was previously suggested to cause mRNA downregulation via U1 snRNP-mediated repression of pre-mRNA 3′ end processing,^43,44^ but direct evidence has not been obtained. In contrast, our model predicts that this SNP should induce mRNA degradation by the exosome through the NRDC mechanism. To test this, we created reporters by inserting the WT or SNP-containing *F9* 3′ UTR sequences, including the PAS, downstream of the eGFP coding sequence (Fig. 5A). When we expressed these reporters in HEK293T cells, we observed a 2-fold decrease in the mRNA level of the SNP-containing Mut compared to that of the WT (p = 0.01, unpaired t-test) (Fig. 5B). As reported previously,^44^ the effect of this mutation was completely reversed when U1 snRNP was inhibited using a U1 antisense morpholino oligo (AMO) (4.5-fold increase, p = 0.0002, unpaired t-test), which blocks U1 snRNP-RNA interactions (Fig. 5C). It is noted that the *F9* WT reporter mRNA level also increased following U1 AMO treatment, albeit much less than that observed for the Mut (2.4-fold increase, p = 0.014, unpaired t-test) (Fig. 5C). This result strongly suggests that the SNP-induced reduction in *F9* mRNA levels is mediated by U1 snRNP. To test if the RNA exosome was involved, we expressed these reporters in the mock (DMSO) or dTAG-treated EXOSC3 degron cell line. Strikingly, dTAG-induced EXOSC3 depletion restored the SNP-containing reporter mRNA levels to that of the WT reporter (2.2-fold increase, p = 0.0203, unpaired t-test), but had little effect on the WT *F9* reporter (Fig. 5D). These results strongly suggest that this SNP-created 5′ ss in the *F9* 3′ UTR suppresses its mRNA levels by promoting exosome-mediated mRNA degradation.

**Figure 5.**
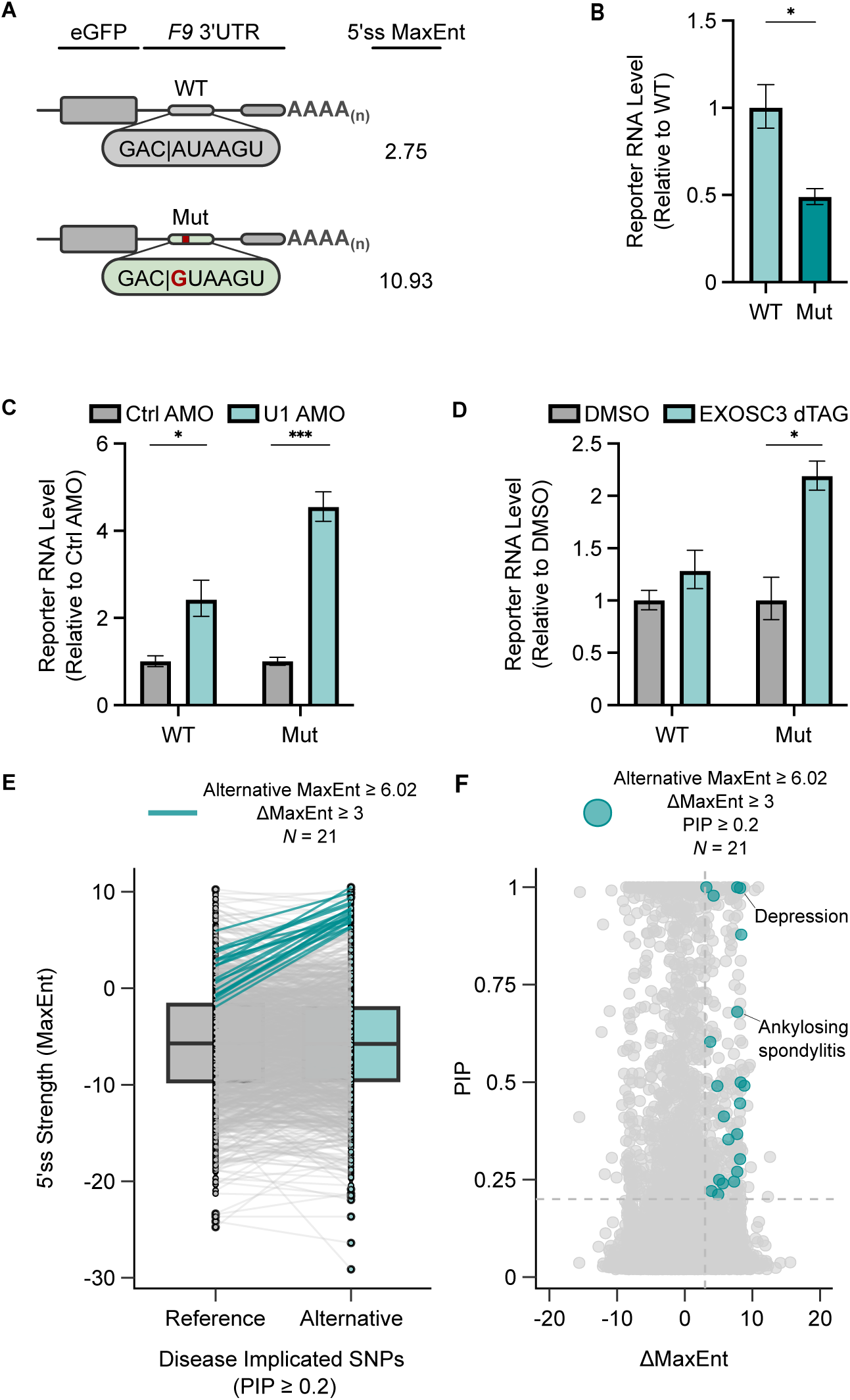
SNP-activated U1 snRNP-dependent mRNA decay in human diseases. **A**, Schematic depicting the eGFP-F9 reporters used in **B** – **D**. Each reporter contains the eGFP coding sequence followed by the wild-type (WT) or patient mutant (Mut) 3′ UTR from the *F9* gene, including the poly(A) site and downstream sequence. The 9 nucleotides shown were used to compute the 5′ ss strength by MaxEnt. The A > G SNP within the *F9* Mut reporter is shown in red. **B**, RT-qPCR analysis of oligo d(T)-primed cDNA following transfection of each reporter into HEK293T cells. Each reporter’s mRNA level was normalized to expression of a co-transfected firefly luciferase control. Expression of the *F9* Mut reporter was then calculated relative to the *F9* WT reporter. Statistical analysis from *n* = 3 independent samples was calculated using an unpaired t-test. **C** & **D**, RT-qPCR analysis of oligo d(T)-primed cDNA following overexpression of each reporter and inhibition of U1 snRNP using U1 AMO (**C**) or dTAG-induced depletion of EXOSC3 (**D**). The reporter mRNA levels were first normalized to the expression of a co-transfected firefly luciferase control plasmid. Then for each reporter, the mRNA level in the U1 AMO condition (**C**) or the EXOSC3 dTAG condition (**D**) was calculated relative to the control condition (Ctrl AMO in **C**, DMSO in **D**). Statistical analysis from *n* = 3 independent samples was calculated using unpaired t-tests. **E**, Boxplot with paired points depicting the 5′ ss strength of all reference and alternative allele variants with a PIP score ≥ 0.2. Lines connect each reference and variant SNP MaxEnt score. Variants marked with a teal line increase the reference allele 5′ ss strength by at least 3 MaxEnt units to an alternative allele 5′ ss strength ≥ 6.02 (1 IQR below the median 5′ ss strength across all 5′ splice sites). **F**, Scatterplot depicting the change in MaxEnt score (ΛλMaxEnt) and the PIP score for all analyzed variants. Variants marked with a teal circle have a PIP score ≥ 0.2 and increase the reference allele 5′ ss strength by at least 3 MaxEnt units to an alternative allele 5′ ss strength ≥ 6.02 (1 IQR below the median 5′ ss strength across all 5′ splice sites). The variant-associated disease or trait is marked for two SNPs.

Similarly, a SNP in the 3′ UTR of the *p14/LAMTOR2* gene causes p14 deficiency, a primary immunodeficiency syndrome.^45,46^ This SNP creates a 5′ ss that leads to U1 snRNP-dependent downregulation of *p14* mRNAs through an unclear mechanism.^45^ We also generated reporters that included the *p14* coding sequence and WT or SNP-containing 3′ UTR downstream and observed a 6.25-fold decrease in the mRNA levels of the SNP-containing reporter compared to the WT (p < 0.0001, unpaired t-test) (Fig. S4A). This reduction was mostly rescued upon EXOSC3 depletion (3.39-fold increase, p < 0.0001, unpaired t-test) (Fig. S4B). By contrast, EXOSC3 depletion had a mild repressive effect on the mRNA levels of the WT *p14* reporter (1.4-fold decrease, p = 0.0252, unpaired t-test) (Fig. S4B). Together our analyses of the SNPs within the *F9* and *p14* genes strongly suggest that these SNPs create novel 5′ ss in the 3′ UTR and induce aberrant mRNA degradation via the NRDC pathway. Thus, we will refer to this type of SNPs as NRDC-activating SNPs. Furthermore, our data provided evidence that exosome depletion or inhibition may be a therapeutic strategy for diseases caused by NRDC-activating SNPs.

To begin identifying additional NRDC-activating SNPs that may contribute to human diseases, we extracted 7,848 fine-mapped SNPs within the annotated 3′ UTRs of all human genes from the CAUSALdb^47^ and UK Biobank^48^ databases and identified those that create novel 5′ splice sites. We selected 106 SNPs that increased the 5′ ss MaxEnt scores of the surrounding sequence by at least 3 units to 6.02 or higher, which is 1 interquartile range (IQR) below the median 5′ ss strength across all annotated introns (mean: 8.66, IQR: 2.64). Next we filtered these SNPs based on their posterior inclusion probability (PIP) scores, which is a measurement of the probability that a variant is a causal factor for a disease or trait.^49^ We selected those SNPs whose PIP score was at least 0.2, which has been used in previous studies to select causal variants.^50^ Using these parameters, we identified 21 SNPs that created a novel 5′ ss (Fig. 5E) and were associated with a wide range of diseases/disorders, including ankylosing spondylitis and depression (Fig. 5F, Supplemental Table 1). Similar to the aforementioned SNPs in *F9* and *p14*, we propose that these SNPs could contribute to diseases by promoting aberrant mRNA degradation via the NRDC mechanism. Using a similar strategy, we also identified 8 potential NRDC-inactivating SNPs. These SNPs are found in naturally occurring 5′ splice sites in 3′ UTRs (MaxEnt ≥ 6.02), indicating that these mRNAs may be naturally unstable due to the NRDC mechanism. These SNPs reduced the 5′ ss strength scores of the surrounding sequences by at least 3 units (Fig. S4C-D, Table S1), which we predict would lead to stabilization of these mRNAs. These NRDC-inactivating SNPs included a variant strongly associated with schizophrenia (Fig. S4D, Table S1). Taken together, our results identified the first NRDC, the 5′ ss-poly(A) junction combination, that induces RNA degradation by the exosome and demonstrated that genetic variations, including SNPs, can contribute to human diseases by disrupting the NRDC pathway and causing aberrant RNA degradation.

## DISCUSSION

A central outstanding question in the RNA degradation field has been how the substrate specificity of the RNA degradation enzymes/machinery is determined. Previous studies have identified several individual RNA sequence motifs that stimulate RNA degradation, such as AU-rich elements in 3′ UTRs,^51^ a “determinant of selective removal” (DSR) sequence in meiotic transcripts,^52^ and short sequence motifs in cryptic unstable transcripts (CUTs).^5^ However, such destabilizing sequence features have not been identified for most unstable RNAs. In this study, we demonstrate that the combination of a 5′ ss and a poly(A) junction, but not either sequence alone, drives exosome-mediated RNA degradation. Given its combinatorial nature, we refer to the 5′ ss-poly(A) junction combination as a “nuclear RNA degradation code” (NRDC) (Fig. 6). This mechanism is fundamentally distinct from the single motif-induced degradation mechanisms in that neither of the individual RNA features per se is destabilizing. 5′ splice sites are typically found in unspliced precursor RNAs whereas poly(A) junctions are found in processed RNAs. The unusual combination of these sequences on the same RNA molecule, as occurs within IPA transcripts and misprocessed RNAs, serves as the degradation mark (Fig. 6). Thus, this mechanism allows for cells to monitor the RNA processing status of all RNA molecules using the processing machinery themselves. We speculate that the concept of the RNA degradation code, i.e. a combination of multiple RNA features serving as the degradation mark, may be broadly applicable for other groups of exosome targets.

**Figure 6.**
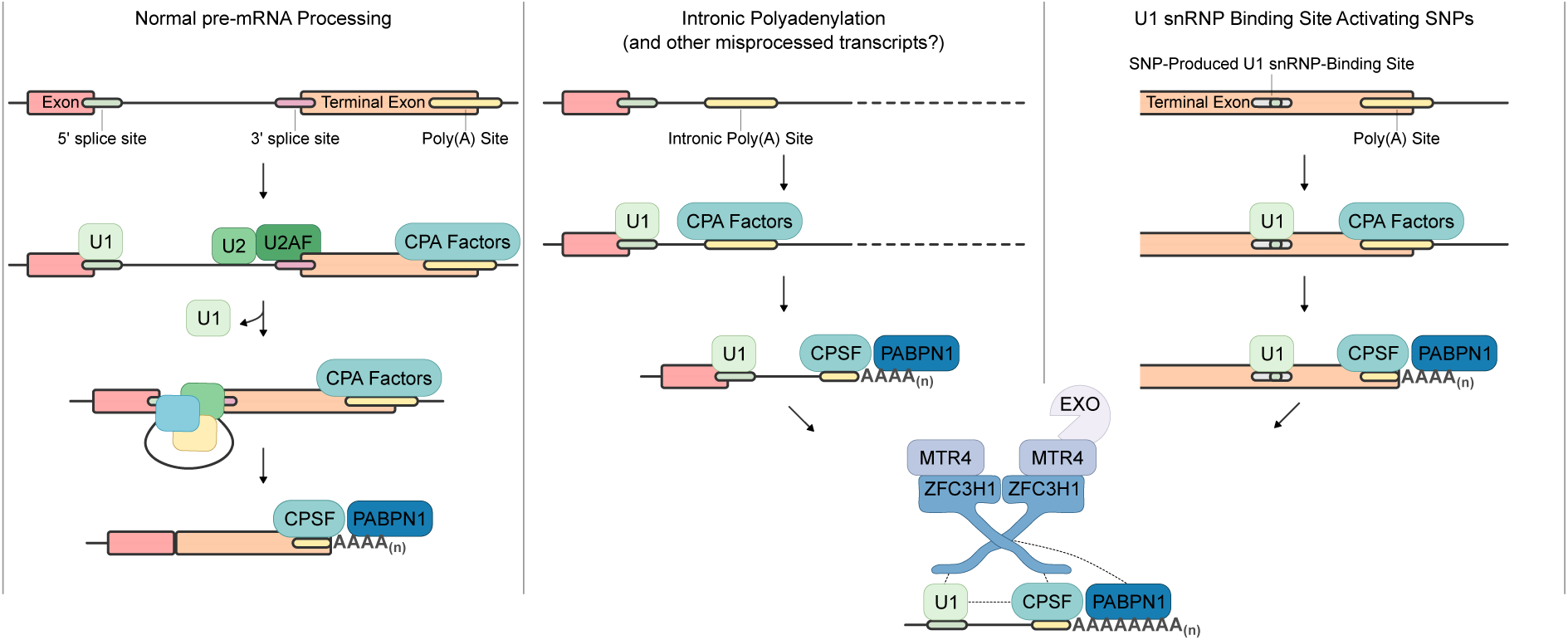
Nuclear RNA Degradation Code. A proposed model depicting how the Nuclear RNA Degradation Code is selectively engaged when intronic polyadenylation occurs or when SNPs provide novel 5′ splice sites in 3′ UTRs, but not during normal pre-mRNA processing. Left: During normal pre-mRNA processing, the 5′ ss is recognized by U1 snRNP, and the 3′ ss and the upstream branch point sequence is recognized by U2AF and U2 snRNP, respectively. During the splicing reaction, U1 snRNP departs from the pre-mRNA and the 5′ ss is removed once splicing is completed. The CPA factors recognize the poly(A) site within the terminal exon and cleave and polyadenylate the pre-mRNA, completing pre-mRNA processing. Middle: When intronic polyadenylation occurs, U1 snRNP remains aberrantly bound at the 5′ ss after cleavage and polyadenylation has occurred in the intron. Right: Similarly, SNPs in the 3′ UTR that create a novel 5′ ss near a poly(A) site result in U1 snRNP binding near a poly(A) site in the terminal exon. Middle/Right: In both cases, the combination of a 5′ ss and a nearby poly(A) junction, bound by U1 snRNP and CPA factors respectively. We propose that this leads to the cooperative recruitment of a PAXT core dimer and degradation by the nuclear RNA exosome via the Nuclear RNA Degradation Code mechanism.

Mechanistically, the 5′ ss and the poly(A) junction are recognized by U1 snRNP and CPA factors, which in turn synergistically recruit the core PAXT components ZFC3H1 and MTR4. Given that U1 snRNP and CPSF bind to a similar region in ZFC3H1 (Fig. 4B-C), and that ZFC3H1 is predicted to form a dimer,^41^ it is possible that U1 snRNP and CPSF cooperatively recruit a ZFC3H1/MTR4 dimer, which in turn recruits the exosome (Fig. 6). Importantly, we showed that PABPN1 is not sufficient to recruit PAXT and instead a poly(A) junction containing a wild-type upstream poly(A) site sequence is required (Fig. 3). Such requirement for a poly(A) junction, instead of simply a poly(A) tail, may prevent spurious degradation of pre-mRNAs that contain an A-rich intronic sequence downstream of the 5′ ss, further increasing specificity. Given the complexity of the interactions between ZFC3H1, U1 snRNP, and CPA factors, crosslinking and structural studies are needed in the future to fully dissect the NRDC recognition complex.

NRDC-mediated degradation is likely subject to regulation. For example, many IPA transcripts are stable and readily detected under steady-state conditions. Our analyses of these transcripts revealed that their upstream 5′ ss is generally weaker than those of IPA transcripts targeted for PAXT-dependent exosomal degradation (Fig. 1E), suggesting that the affinity of U1 snRNP binding to the upstream 5′ ss is an important determinant of the efficiency of NRDC-mediated RNA degradation. Further, it is well known that there are stable polyadenylated mRNAs that have retained introns. Some of them are exported into the cytoplasm and others are sequestered in the nucleus; the introns of the latter group are called “detained introns.”^53,54^ How do they remain stable? Previous studies have provided evidence that, although these retained introns are not spliced, spliceosomes do assemble on them.^55^ Therefore, the U1 snRNP complexes on these transcripts are most likely engaged with U2AF and other spliceosome components and, thus, cannot associate with CPA factors to synergistically recruit PAXT. For future studies, it will be important to determine whether and how the distance and the RNA sequences between the 5′ ss and the poly(A) junction can modulate the degradation efficiency.

In addition to its role in transcript surveillance, the NRDC may also play an important role in gene evolution. For example, many lncRNAs reside in the nucleus and previous studies provided evidence that their nuclear retention is mediated by their interaction with U1 snRNP.^31,37^ Some of these nuclear lncRNAs are highly abundant and stable, including *MALAT1* and *NEAT1*. Interestingly, both *MALAT1* and the long isoform of *NEAT1*, *NEAT1_2*, have non-polyadenylated 3′ ends.^32^ Our NRDC model predicts that both *MALAT1* and *NEAT1* would be targeted for exosomal degradation if they had poly(A) tails. Indeed, the 3′ end of the short isoform of *NEAT1*, *NEAT1_1*, is formed through the canonical CPA process,^56^ and our data showed that it is targeted by PAXT-mediated exosomal degradation (Fig. S5A-C). Thus, it is likely that both *MALAT1* and *NEAT1_2* have evolved non-polyadenylated 3′ ends to evade degradation via the NRDC mechanism.

Our study linked the NRDC mechanism to human diseases. Two SNPs in the *F9* and *p14* genes have been shown to cause severe hemophilia B and p14 deficiency respectively. These SNPs create a novel 5′ ss in the 3′ UTR and cause significant decreases in the mRNA and protein levels of these genes.^43–45^ Previous studies proposed that these SNP-created 5′ splice sites repress mRNA 3′ processing,^44,45^ but direct evidence has been lacking. Our data demonstrated that the SNP-induced downregulation of mRNA levels in *F9* and *p14* genes could be reversed by EXOSC3 depletion, strongly suggesting that these SNPs primarily act to induce mRNA degradation rather than to block mRNA 3′ processing (Fig. 5D and Fig. S4B). Our analyses of several public databases of genetic variations have identified over 20 additional SNPs that may contribute to disease through a similar NRDC mechanism. Our initial survey serves as a proof-of-principle study and is by no means comprehensive. For example, none of the databases that we searched contained the disease-causing SNPs reported for *F9* and *p14*. Thus, more extensive analyses are needed in the future to uncover NRDC-activating genetic variations at the genome-wide level. Finally, our data showed that the SNP-induced decrease in *F9* and *p14* mRNA levels could be fully reversed to near wild-type levels, raising the potential for use of exosome inhibition as a therapeutic strategy for these diseases. Alternatively, antisense oligonucleotides that hybridize to the SNP-containing regions may prevent U1 snRNP binding and in turn prevent RNA degradation. Taken together, our study highlights how mechanistic analyses of a fundamental pathway, such as the NRDC, can guide future efforts in therapeutic development.

### Data Availability

PAS-seq datasets have been deposited into the GEO under accession number GSE267857.

## Acknowledgments

We would like to thank Drs. Shalini Sharma, Doug Black, Bert Semler, and François Bachand for providing reagents; Ivan Marazzi, Feng Qiao, and Johannes Linder for their comments on the manuscript. We wish to acknowledge the support of the Chao Family Comprehensive Cancer Center Shared Resource Genomics High-Throughput Facility, supported by the National Cancer Institute of the National Institutes of Health under award number P30CA062203. We would like to thank Seung-Ah Yoon from the UCI Genomics High-Throughput Facility, and Adeela Syed from the UCI Optical Biology Core Facility for technical support. This study was supported by the following grants: R35GM149294 (Y.S.), NIH Director’s New Innovator Award, DP2GM149554 (H.Y. and F.D.). L.T. is supported by the Hewitt Foundation Postdoctoral Fellowship.

## Author Contributions

Conceptualization: L.V.S. and Y.S.; Investigation: L.V.S., L.Liu, X.Z., Y.Y.; Formal Analysis: L.V.S., X.Z., A.Y., W.L., Y.S.; Methodology: L.V.S., H.Y., L.T.; Resources: H.Y.; Validation: L.V.S., S.L., L.Liu, L.T., M.C.V.; Writing – Original Draft: L.V.S.; Writing – Review & Editing: L.V.S., Y.S.; Funding Acquisition: Y.S.; Supervision: Y.S.

## METHODS

### Cell Lines and Cell Culture Conditions

#### Cell Culture Conditions

Wild-type and edited HEK293T cell lines were maintained in DMEM (Gibco) supplemented with 10% FBS (Sigma). Cells were grown at 37°C with 5% CO_2_. Cell lines were routinely monitored for mycoplasma contamination using the MycoAlert Mycoplasma Detection Kit (Lonza).

#### Generation of FLAG-dTAG Edited HEK293T Cell Lines

The C-terminal dTAG-FLAG knockins of EXOSC3 and the N-terminal FLAG-dTAG knockin of ZFC3H1 in HEK293T cells were achieved using CRISPR/Cas9 and the microhomology end joining (MMEJ) approach as described in,^57^ except that we used Lipofectamine 3000 to perform the transfections. To construct the donor vector, we modified pCRIS-PITCHv2-dTAG-BSD (BRD4) and pCRIS-PITCHv2-BSD-dTAG (BRD4) to replace the HA tag with a 3X-FLAG tag and then replaced the microhomology sequences for EXOSC3 or ZFC3H1 using In Fusion cloning (Takara). pCRIS-PITChv2-dTAG-BSD (BRD4) and pCRIS-PITCHv2-BSD-dTAG (BRD4) were gifts from James Bradner & Behnam Nabet (Addgene plasmid # 91795; https://www.addgene.org/91795/ and Addgene plasmid # 91792; https://www.addgene.org/91792/).^58^ We used the online tool CHOPCHOP^59^ to select a specific and efficient sgRNA. We screened and validated single colonies using genomic DNA PCR followed by Sanger Sequencing where possible and by western blotting for all cell lines. Homozygous edited colonies were selected and used in experiments. For ZFC3H1, only the largest isoform exhibited dTAG-dependent degradation.

### RNAi

#### Lentiviral Knockdown

Lentiviral particles were prepared by co-transfecting HEK293T cells with the following plasmids: pLKO: shRNA vector, PMD2.G, and psPAX2. The virus-containing cell culture media was collected 24 and 48 hours after transfection and passed through a 45 µM filter. The titer was measured using a qPCR-based kit (AbmGood). HEK293T cells were transduced at an MOI of 10 with lentivirus expressing one or two different shRNAs targeting MTR4, ZFC3H1, PABPN1 or a control scramble sequence. Polybrene (8 µg/mL) was included during transduction. 24 hours after transduction, puromycin was added to the media at a concentration of 1.25 µg/mL. 4 days after transduction, cells were harvested in Trizol for RNA purification or directly lysed in 1X SDS loading dye for western blotting.

#### siRNA Knockdown

An EXOSC3-targeting siRNA (IDT) or a pool of control siRNAs (Sigma) was transfected into cells using Lipofectamine RNAiMAX at a final siRNA concentration of 20 nM. 48 hours later, the transfection was repeated. 24 hours later, cells were harvested in Trizol for RNA purification or directly lysed in 1X SDS loading dye for western blotting.

### PAS-seq

#### Library Preparation

PAS-seq libraries were prepared as previously described^25^ with the following minimal modifications. Purified mRNA was fragmented at 94°C for 3 minutes using the NEBNext Magnesium RNA fragmentation module. In addition, the libraries were resolved on a 2.5% agarose gel and the region between 200-300 basepairs was purified by gel extraction and sequenced on the Illumina NovaSeq6000 platform. For all samples, a small amount of mouse RNA was spiked in but was not used in data processing. For the spike-ins, an extra control or treatment well was prepared and the cells in this extra well were counted and recorded. 1 µg total RNA from each sample was mixed with up to 10% by mass (100 ng) mouse total RNA (purified from mouse brain tissue) prior to library preparation. The spike-in RNA mass was calculated relative to the measured cell count for each sample. For example, if there were 1 x 10^6^ control cells and 0.8 x 10^6^ treated cells, then 100 ng mouse spike-in RNA would be added to 1 µg of control total RNA and 80 ng mouse spike-in RNA would be added to 1 µg of treated total RNA. As such, the level of spike-in RNA could be used to normalize sequencing reads to the original cell number.

#### PAS-seq Data Processing

PAS-seq data processing was conducted as previously described with few modifications.^60^ Briefly, we used Cutadapt (v2.10)^61^ to produce trimmed PAS-seq reads by: 1) trimming the 6 nucleotide linker sequence, 2) trimming the poly(A) tail sequence, and 3) removing any untrimmed reads. Trimmed reads were then mapped to a concatenated hg19 and mm9 genome using STAR (v2.7.3a)^62^ with the --alignEndsType EndToEnd parameter. The mapped reads were converted from a bam file to a bed file using bedtools (v2.29.2)^63^. Using a custom python script, reads were removed as potential internal priming events if they mapped to a genomic region where 6 consecutive A’s or 7 A’s out of 10 nucleotides were observed in the 10 nucleotides downstream of the read. The remaining reads were converted to a bam file and used to generate bigwig files using bedtools. The 3’ ends of the reads were extracted from the bed files using bedtools flank and the initial read counts for each poly(A) site were calculated using bedtools coverage with a master file of all annotated poly(A) sites.

#### Differential Isoform Expression Analysis

To detect differentially expressed transcripts using PAS-Seq data, we treated counts mapping to each poly(A) site region as a “gene” and analyzed differential gene expression using the edgeR (v3.40.1) topTags function. We considered a transcript to be differentially expressed if it displayed an FDR ≤ 0.05 and a log_2_FC > 1 or log_2_FC < -1, unless indicated otherwise.

### Molecular Cloning

#### eGFP-5′ ss Reporters

The eGFP-5′ ss reporters contained the FLAG-eGFP coding sequence followed by a 3′ UTR in the pCDNA5 expression vector. In the 5′ ss-containing reporters, the 3′ UTR included a 28 nucleotide sequence from the gene *NXF1* that was previously shown to contain one strong and one weaker 5′ ss.^31^ In the 5′ ss mutant reporters, both 5′ splice sites were mutated by PCR mutagenesis. The polyadenylated reporters contained either the bovine growth hormone (bGH) poly(A) site or the adenovirus major late (L3) poly(A) site downstream of the 5′ ss. The non-polyadenylated reporters contained either: 1) a 164 nucleotide sequence from the 3′ end of *MALAT1* or, 2) a 129 nucleotide region including part of the *H2AC18* 3′UTR and 56 nucleotides of downstream sequence to elicit histone 3′ end processing.

#### RNA Pulldown Constructs

The RNA pulldown constructs contained 3 copies of the MS2 hairpin sequence followed by the 5’ ss sequence 5’-CAG|GTAAGT-3′ and the L3 poly(A) site in the pBluescript vector. The distance between the end of the 5′ ss and the L3 AAUAAA poly(A) signal was 68 nucleotides. In the 5′ ss mutant constructs, the 5′ ss was mutated to 5′-AGA|AGCCAT-3′. In the poly(A) site mutant constructs, the L3 poly(A) signal was mutated from 5′-AAUAAA-3′ to 5′-AA**G**AAA-3′.

#### FLAG-ZFC3H1 Expression Constructs

The FLAG-ZFC3H1 expression constructs were cloned into pCDNA3. The truncations and mutations were produced by PCR mutagenesis. The ZFC3H1 SLiM mutant contained the following mutations: 18-EEGELEDGEI-27 to 18-E**A**G**A**LE**A**G**A**I-27. All constructs contained an N-terminal FLAG tag and the c-myc nuclear localization signal was added to all expression constructs except the full length.

#### eGFP-F9 Reporters

For the eGFP-F9 reporters, the entire *F9* 3′UTR sequence and 597 nucleotides of downstream genomic sequence was cloned downstream of FLAG-eGFP in pCDNA5. The A>G patient mutation 1157 basepairs downstream of the stop codon was introduced by PCR mutagenesis.

### Reporter Assays

For baseline eGFP-5′ss reporter mRNA and protein level measurements, eGFP-5′ ss reporters were co-transfected with the renilla luciferase expressing plasmid pRL-TK at a 10:1 ratio by mass into HEK293T cells (i.e. 250 ng pCDNA5 and 25 ng pRL-TK). 48 hours after transfection, cells were harvested in Trizol for RNA purification or directly lysed in 1X SDS loading buffer for western blotting.

For baseline eGFP-F9 reporter mRNA level measurements, eGFP-F9 reporters were co-transfected with the firefly luciferase expressing plasmid pGL3: promoter at a 10:1 ratio by mass into HEK293T cells. 24 hours after transfection, cells were harvested in Trizol for RNA purification.

To measure the mRNA level of reporters after dTAG-induced depletion of EXOSC3 or ZFC3H1, or U1 inhibition by U1 AMO, eGFP-5′ ss or eGFP-F9 reporters were co-transfected with the firefly luciferase expressing plasmid pGL3: promoter at a 10:1 ratio by mass into HEK293T cells. To deplete EXOSC3 or ZFC3H1, 16 hours after transfection dTAG^v^-1 or an equivalent volume of DMSO was added to cell culture media at a final concentration of 500 nM and cells were harvested 8 hours later.^35^ To inhibit U1 snRNP using U1 AMO, 16 hours after transfection 25 µM U1 AMO or Control AMO was nucleofected into transfected cells and cells were harvested 8 hours later. Cells were harvested in Trizol for RNA purification or directly lysed in 1X SDS loading buffer for western blotting.

### Inhibition of U1 snRNP by AMO

To inhibit U1 snRNP, 25 µM control or U1 AMO (GeneTools, LLC) was delivered into 1x10^6^cells by nucleofection using the Lonza SF Cell Line 4D-Nucleofector^TM^ X Kit using the DH-135 program. Cells were harvested 8 hours later in Trizol for RNA purification.

### RT-qPCR

Total RNA was purified following the Trizol manufacturer instructions. 500 ng-1 µg total RNA was DNase-treated using RQ1 DNase (Promega) followed by addition of RQ1 DNase stop solution and heat inactivation. For experiments using eGFP-5′ ss reporters, the DNase-treated RNA was reverse transcribed using Superscript III (Invitrogen) and random hexamers or oligo d(T)_20_ where indicated. For experiments using eGFP-F9 and *p14* reporters, the DNase-treated RNA was reverse transcribed using Superscript III and oligo d(T)_20_. The resulting cDNA was diluted and used for qPCR using the PowerUp^TM^ SYBR^TM^ Green Master Mix (Applied Biosystems). All qPCR analyses used the co-transfected plasmid to normalize reporter mRNA measurements (renilla luciferase from pRL:TK or firefly luciferase from pGL3: promoter). The delta-delta Ct method was used to analyze qPCR data as described previously.^64^ Normalized expression data was plotted on the y-axis and confidence intervals were calculated to plot error bars.

### RNA FISH and Immunofluorescence

To monitor RNA localization of eGFP-5′ ss reporters, we seeded 56,250 HEK293T cells per well (225,000 cells/mL) of 8-well glass bottom chamber slides (ibidi) coated with 0.1 mg/mL Poly-D-Lysine (Gibco). The next morning, we transfected 125 ng of each eGFP-5′ ss reporter using Lipofectamine 3000. 12 hours later, we fixed cells with 4% paraformaldehyde for 10 minutes. Cells were then washed with 1X PBS and permeabilized with 70% ethanol at -20°C overnight. RNA-FISH probes targeting the eGFP coding sequence were synthesized as described previously.^65^ To hybridize probes to the reporter RNAs, cells were washed once with wash buffer (2X SSC (Invitrogen), 20% deionized formamide (Ambion), and 0.1% Tween-20 (ThermoFisher)) and then incubated overnight at 30°C with 125 ng of RNA-FISH probes in 100 µL hybridization buffer (0.1 g/mL dextran sulfate (Fisher), 0.1 mg/mL E.coli tRNA (Roche), 2 mM Vanadyl ribonucleoside complex (NEB), 2X SSC, 20% deionized formamide, 0.1 % Tween-20). The next day, cells were washed twice with 30°C wash buffer and then incubated for 30 minutes in wash buffer. Cells were post-fixed with 4% paraformaldehyde for 10 minutes and incubated with immunofluorescence primary antibodies for one hour at room temperature in PBS-Tween 20 (PBST). Cells were washed once with PBST and then incubated with immunofluorescence secondary antibodies for 30 minutes at room temperature in PBST. Cells were washed with PBST and stained for DAPI (Invitrogen) using a 1:20,000 dilution for 5 minutes. Finally, cells were mounted with Fluoromount G (Invitrogen) before imaging. Imaging was performed using the Leica DMi8 THUNDER. Ten to twelve images per reporter were taken. The nuclear versus cytoplasmic RNA FISH signal was quantified using ImageJ. All transfected cells were manually selected and the total nuclear FISH intensity was divided by the total cytoplasmic FISH intensity per image.

### *In vitro* Transcription

For RNA pulldowns and polyadenylation assays, 3XMS2-RNA was *in vitro* transcribed using T7 RNA Polymerase (NEB) and free nucleotides were removed using a column (Cytiva). For radiolabeled RNA substrates, [α-^32^P]-UTP (PerkinElmer) was included in the transcription reaction. The RNA was purified by phenol/chloroform extraction followed by ethanol precipitation. For radiolabeled RNAs only, the RNA was further subjected to Urea-PAGE purification and ethanol precipitated. To produce the pre-polyadenylated substrates, the DNA template for transcription was prepared by PCR linearizing the pBluescript constructs at the cleavage site and used for *in vitro* transcription. The resulting “pre-cleaved” RNA was polyadenylated by *E. coli* Poly(A) Polymerase (NEB) and passed through a column (Cytiva) to remove free nucleotides and purified by phenol/chloroform extraction followed by ethanol precipitation.

### *In vitro* Polyadenylation Assay

1 pmol (20 cps) radiolabeled RNA was mixed with 1 µL 0.1 M ATP, 2 µL 1 M creatine phosphate, 1 µL 10 µg/µL yeast tRNA, 4.4 µL HeLa nuclear extract, and H_2_O to a final volume of 10 µL. The reaction was incubated at 30°C for 20 minutes or 150 minutes as indicated. The proteins in the reaction were then digested using Proteinase K and the RNA was purified by phenol/chloroform extraction followed by ethanol precipitation. The RNA was then resolved on an 8% Urea-PAGE gel. The gel was transferred to filter paper, vacuum dried, and used to expose a phosphor screen overnight. The phosphor screen was imaged using a GE Amersham^TM^ Typhoon^TM^.

### MS2-MBP RNA Pulldown

11.25 pmol RNA and 112.5 pmol of MBP-MS2 fusion protein was brought to 50 µL with Buffer D-100 (20 mM HEPES-NaOH pH 7.9, 100 mM NaCl, 1 mM MgCl2, 0.2 mM EDTA) and incubated on ice for 30 minutes. The following was then added: 7.5 µL 0.1 M ATP, 15 µL 1 M creatine phosphate, 7.5 µL 10 µg/µL yeast tRNA, 300 µL HeLa nuclear extract, and H_2_O to a final volume of 750 µL. The reaction was incubated at 30°C for 20 minutes. The reaction was then chilled on ice, centrifuged for 1 minute at 14,000 x rpm (>16,000 x g) at 4°C. The supernatant was mixed with pre-washed amylose beads (55 µL slurry) and rotated for 1 hour at 4°C. The beads were washed three times for 10 minutes per wash with 1 mL wash buffer (20 mM HEPES-NaOH pH 7.9, 100 mM KCl, 4% glycerol, 1 mM DTT) supplemented with detergent (0.05% IGEPAL CA-630). The beads were washed once more for 10 minutes per wash with 1 mL wash buffer without detergent. Bound proteins were eluted twice using 125 µL of wash buffer without detergent supplemented with 20 mM maltose for 10 minutes per elution. The eluted proteins were acetone precipitated overnight at -20°C. The next day, samples were centrifuged for 20 minutes at 14,000 x rpm, pellets were resuspended in 1X SDS loading buffer, heated to 95°C for 7 minutes, and subjected to standard western blotting procedures. All pulldown protein samples were resolved on 4-20% pre-cast SDS-PAGE gels (BioRad) and subjected to a “standard” eBlot transfer.

### FLAG-Immunoprecipitation

2.2x10^6^ HEK293T cells were seeded per 10-cm dish. The next day, pCDNA3: ZFC3H1 expression plasmids (Full length or truncated) were transfected. The transfected DNA amount was titrated in an effort to achieve approximately equal levels of each construct. Accordingly, we transfected the following amounts: 8 µg empty vector, 8 µg full-length ZFC3H1, 10 µg N-term, 24 µg C-term, 8 µg N-term 1SLiM, 8 µg 50-599, 16 µg 600-990. Two days after transfection, cells were scraped into 5 mL cold 1X PBS, centrifuged at 100 x g for 3 mins, and the cell pellets were flash frozen on dry ice and stored at -80°C overnight. The next day, cells were resuspended in 1 mL cold lysis buffer per plate (20 mM HEPES-NaOH pH 7.9, 150 mM NaCl, 2 mM EDTA, 0.1% Triton X-100, 1 mM PMSF added fresh, 1X Thermo Scientific Halt^TM^ Protease Inhibitor Cocktail added fresh). While on ice, each sample was sonicated with a microtip at amplitude 1, for 3 seconds on, 10 seconds off, repeated 6 times total. The lysate was then cleared by centrifugation at 14,000 x rpm for 30 minutes at 4°C and an aliquot was removed for input. During this time, 40 µL of Anti-FLAG M2 agarose beads slurry (Sigma) was washed 3 times with 1 mL lysis buffer per wash. For + RNase A samples, 2 µL RNase A (20 µg RNase A/IP) was added to the supernatant and mixed by pipetting. The supernatant was then added to the pre-washed beads and rotated at 4°C for 2 hours. The beads were washed 4 times with 1 mL lysis buffer for 10 minutes per wash. The bound proteins were then eluted three times with 100 µL lysis buffer supplemented with 3XFLAG peptide, 10 minutes per elution. The eluted proteins were acetone precipitated overnight at -20°C. The next day, the samples were centrifuged at 14,000 x rpm for 20 minutes and the pellets were resuspended in 1X SDS loading buffer and heated to 95°C for 7 minutes prior to western blotting. For the FLAG western blot, the samples were resolved on a 4-20% pre-cast gel (BioRad) and subjected to a “long” eBlot transfer. For all other proteins, the samples were resolved on a hand-poured 10% SDS-PAGE gel and subjected to a “standard” eBlot transfer.

### SNP Analyses

#### Selection of SNPs

To analyze all potentially causal variants, all fine-mapped, single-nucleotide variants within 3′ UTRs with a PIP score of at least 0.02 were extracted from the CAUSALdb and UK Biobank databases. SNPs that overlapped with 3′ UTRs were identified and downloaded as described in a previous study.^66^

#### Prediction of Nuclear RNA Degradation Code Activating and Inactivating SNPs

To compute the 5′ ss strength of the reference sequence and the variant sequence, we extracted 25 nucleotides of flanking genomic sequence upstream and downstream of the variant position using bedtools slop and bedtools getfasta. We then combined the upstream and downstream genomic sequence with the reference nucleotide or the variant nucleotide in the middle position. We then extracted 9 nucleotide “5′ ss” sequences, wherein the reference or variant nucleotide was located in every possible position (9 permutations for reference and variant). We calculated the 5′ ss strength (MaxEnt) for all sequences.^30^ Finally, we selected the maximum 5′ ss strength score calculated for each reference and variant and calculated the difference between these values. We identified potentially disease-causing, nuclear RNA degradation code-activating variants by filtering for variants with a PIP score of at least 0.2 that increased the reference maximum 5′ ss strength by at least 3 MaxEnt units to 6.02 or higher (1 IQR below the median 5′ ss strength: 8.66). We identified potentially disease-causing, nuclear RNA degradation code-inactivating variants by filtering for variants with a PIP score of at least 0.2 that decreased the reference maximum 5′ ss strength of at least 6.02 (1 IQR below the median 5′ ss strength: 8.66) by at least 3 MaxEnt units.

### Statistics

All statistics were performed using the GraphPad Prism 10.2.1 software. Statistical tests used are listed in the figure legends. Where applicable, unlabeled indicates p-value > 0.05, * indicates p-value ≤ 0.05, ** indicates p-value ≤ 0.01, *** indicates p-value ≤ 0.001, and **** indicates p-value ≤ 0.0001. For all boxplots, the Tukey method was used for error bars. All t-tests and Mann-Whitney tests were two-tailed.

### Antibodies used in this study

#### Primary antibodies

Anti-FLAG mouse (Sigma-Aldrich, F1804-200UG)

Anti-beta Tubulin (Invitrogen, PA1-16947)

Anti-GAPDH (Santa Cruz, sc-365062)

Anti-ZFC3H1 (CCDC131) (Bethyl, A301-456A)

Anti-MTR4 (Abcam, AB70551)

Anti-PABPN1 (Bethyl, A303-524A)

Anti-ZCCHC8 (Bethyl, A301-806A)

Anti-EXOSC3 (Proteintech, 15062-1-AP)

Anti-U1-70K (Millipore, clone 964.1, 05-1588)

Anti-U1A (Santa Cruz, sc-101149)

Anti-U1C (Sigma, SAB4200188-200UL)

Anti-ARS2 (Bethyl, A304-550A)

Anti-CPSF100 (Bethyl, A301-581A)

Anti-CPSF73 (Bethyl, A301-091A) Anti-CPSF30 (Bethyl, A301-585A)

Anti-FLAG rabbit (Cell Signaling, 14793S)

Anti-SC-35* (SRRM2) (Abcam, ab11826)

*Recently, a study reported that anti-SC35, a widely used marker of nuclear speckles, recognizes SRRM2^67^

#### Secondary antibodies

IRDye 680RD Goat Anti-Rabbit IgG (Licor 926-68071)

IRDye 800CW Goat Anti-Mouse IgG (Licor 926-32210)

Goat Anti-Rabbit IgG HRP-conjugate (Millipore 12-348)

Goat Anti-Mouse IgG HRP-conjugate (Millipore 12-349)

Rabbit Anti-Rat IgG HRP-conjugate (Invitrogen PA1-28573)

### Primers used in this study

eGFP-qPCR-F: 5′-CAAGGACGACGGCAACTACAAG-3′

eGFP-qPCR-R: 5′-CTGCTTGTCGGCCATGATATAGAC-3′

Firefly-qPCR-F: 5′-CTCACTGAGACTACATCAGC-3′

Firefly-qPCR-R: 5′-TCCAGATCCACAACCTTCGC-3′

Renilla-qPCR-F: 5′-AGAGAAAGGTGAAGTTCGTCG-3′

Renilla-qPCR-R: 5′-TCTTGGCACCTTCAACAATAGC-3′

F9-qPCR-F: 5′-TAAGTATCATGTCTCCTTTAACTAGCATACC-3′

F9-qPCR-R: 5′-GTCAGTTGGTCAGCAACTCTCTAG-3′

p14-qPCR-F: 5′-CGGTACTGTTGGTAAAGCCACC-3′

p14-qPCR-R: 5′-CAGAGTAGGCCAGCAGTGATCC-3′

### Additional oligos used in this study

EXOSC3 C-terminal knockin sgRNA: 5′-aaaacagatcttctccagat-3′

ZFC3H1 N-terminal knockin sgRNA: 5′-ggagtatctgcggtcgccat-3′

Ctrl siRNA: (Sigma-Aldrich SIC001) EXOSC3 siRNA F (IDT): 5′-rCrArCrGrCrArCrArGrUrArCrUrArGrGrUrCrATT-3′

EXOSC3 siRNA R (IDT): 5′-rUrGrArCrCrUrArGrUrArCrUrGrUrGrCrGrUrGTT-3′

Scramble shRNA F:5′-ccggtcctaaggttaagtcgccctcgctcgagcgagggcgacttaaccttaggtttttg-3′

Scramble shRNA R:5′-aattcaaaaacctaaggttaagtcgccctcgctcgagcgagggcgacttaaccttagga-3′

ZFC3H1 shRNA 1F: 5′-CCGGgtaaggcagacaagcgttattCTCGAGaataacgcttgtctgccttacTTTTTG-3′

ZFC3H1 shRNA 1R: 5′-AATTCAAAAAgtaaggcagacaagcgttattCTCGAGaataacgcttgtctgccttac-3′

ZFC3H1 shRNA 2F: 5′-CCGGgactgatgacatcgctaatttCTCGAGaaattagcgatgtcatcagtcTTTTTG-3′

ZFC3H1 shRNA 2R: 5′-AATTCAAAAAgactgatgacatcgctaatttCTCGAGaaattagcgatgtcatcagtc-3′

MTR4 shRNA 1F: 5′-CCGGagaagttgcttgggttatattCTCGAGaatataacccaagcaacttctTTTTTG-3′

MTR4 shRNA 1R: 5′-AATTCAAAAAagaagttgcttgggttatattCTCGAGaatataacccaagcaacttct-3′

PABPN1 shRNA 1F: 5′-CCGGggagctggaagctatcaaagcCTCGAGgctttgatagcttccagctccTTTTTG-3′

PABPN1 shRNA 1R: 5′-AATTCAAAAAggagctggaagctatcaaagcCTCGAGgctttgatagcttccagctcc-3′

Ctrl AMO: 5′-CCTCTTACCTCAGTTACAATTTATA-3′

U1 AMO: 5′-GGTATCTCCCCTGCCAGGTAAGTAT-3′

## SUPPLEMENTARY METHODS

### U1 snRNA Targeting of 5′ ss mutant reporter

To direct U1 snRNP to bind the 5′ ss mutant bGH poly(A) site reporter, HEK293T cells were co-transfected in a 12-well plate with 250 ng eGFP-5′ss mutant bGH poly(A) site reporter, 250 ng pNS6: U1 snRNA complementary to the 5′ ss mutant or a control sequence, and 25 ng pGL3: promoter as a co-transfection control. 48 hours after transfection, cells were harvested in Trizol for RNA purification. RT-qPCR was performed as described above using cDNA reverse transcribed with oligo d(T)_20_ normalized to firefly luciferase expression to monitor the mRNA levels.

### Reporter Assay Protein Level Quantification

To quantify differences in eGFP-5′ ss reporter protein levels (bGH or *MALAT1* 3′ end), protein lysates were subjected to western blotting using IR dye-conjugated secondary antibodies (Licor). The membranes were imaged using a GE Amersham^TM^ Typhoon^TM^ and quantified using ImageQuant TL software.

### Molecular Cloning

#### p14 Reporters

For the *p14* reporters, the *p14* coding sequence, terminal intron, 3′ UTR, and 317 nucleotides of downstream genomic sequence was cloned into the pGL3: promoter backbone by replacing firefly luciferase. The C>A patient mutation 23 basepairs downstream of the stop codon was introduced by PCR mutagenesis.

### Reporter Assay

#### p14 Reporters

For baseline *p14* reporter mRNA level measurements, *p14* reporters were co-transfected with the firefly luciferase expressing plasmid pGL3: promoter at a 10:1 ratio by mass into HEK293T cells. 24 hours after transfection, cells were harvested in Trizol for RNA purification. RT-qPCR was performed as described above using cDNA reverse transcribed with oligo d(T)_20_ normalized to firefly luciferase expression to monitor the mRNA levels.

To measure the mRNA level of *p14* reporters after dTAG-induced depletion of EXOSC3, *p14* reporters were co-transfected with the firefly luciferase expressing plasmid pGL3: promoter at a 10:1 ratio by mass into HEK293T cells. To deplete EXOSC3, 16 hours after transfection dTAG^v^-1 or an equivalent volume of DMSO was added to cell culture media at a final concentration of 500 nM and cells were harvested 8 hours later.^35^ Cells were harvested in Trizol for RNA purification or directly lysed in 1X SDS loading buffer for western blotting. RT-qPCR was performed as described above using cDNA reverse transcribed with oligo d(T)_20_ normalized to firefly luciferase expression to monitor the mRNA levels.

## SUPPLEMENTAL FIGURES

**Supplemental Figure 1.**
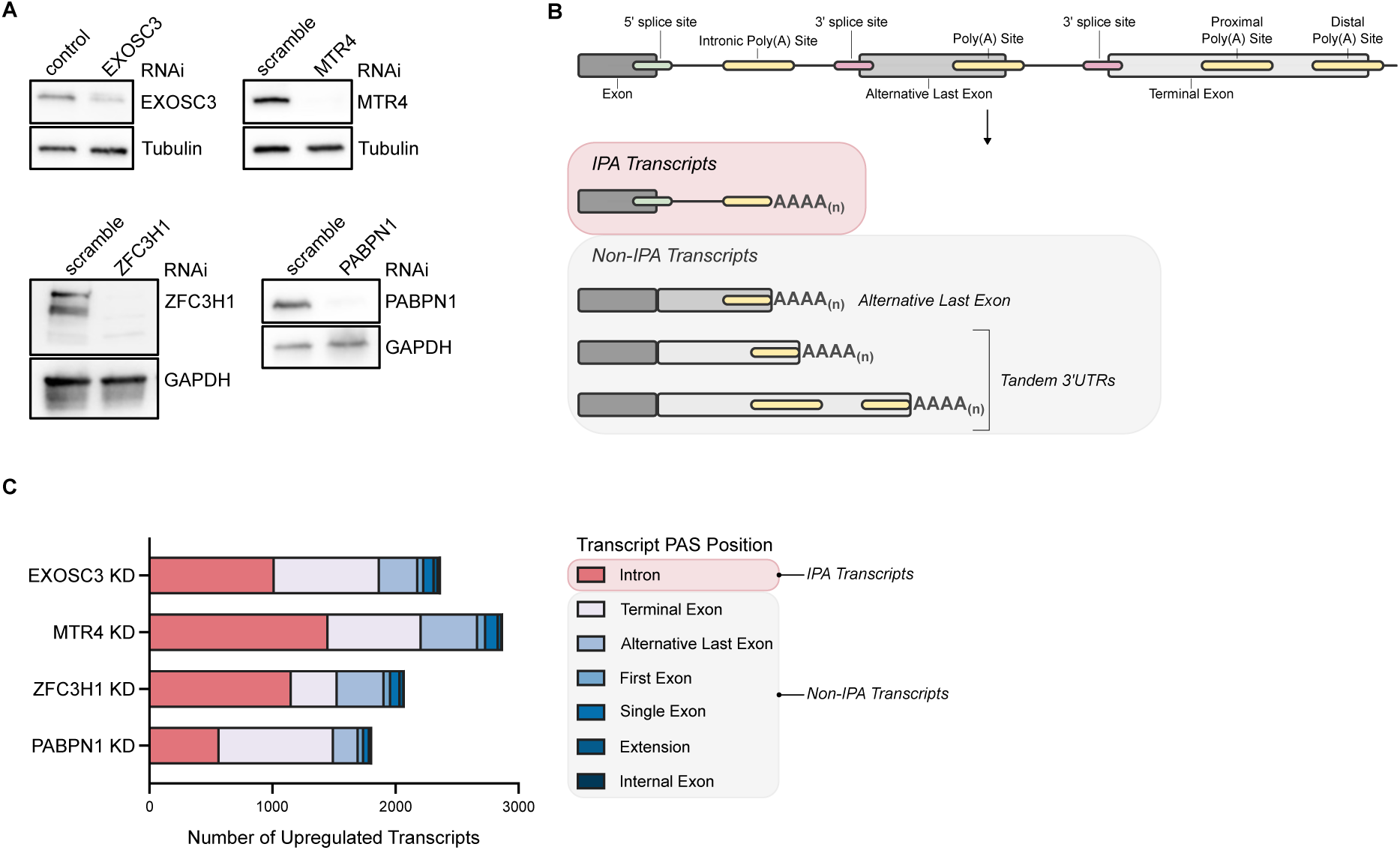
Characterization of gene expression changes following depletion of PAXT components or the RNA exosome. **A**, Western blot analysis of knockdown samples prior to PAS-seq to confirm depletion of EXOSC3, MTR4, ZFC3H1, or PABPN1. Tubulin and GAPDH were used as loading controls. **B**, Schematic depicting the synthesis of IPA transcripts and three common examples of non-IPA transcripts. IPA transcripts are produced when a pre-mRNA is cleaved and polyadenylated within the intron, leaving an intact 5′ ss upstream of the poly(A) tail. For non-IPA transcripts, cleavage and polyadenylation may occur within the terminal exon or within an alternative last exon following alternative splicing. **C**, Stacked barplots depicting the frequency of PAS position across all upregulated transcripts (log_2_FC > 1, FDR ≤ 0.05) in HEK293T cells depleted of EXOSC3, MTR4, ZFC3H1, or PABPN1.

**Supplemental Figure 2.**
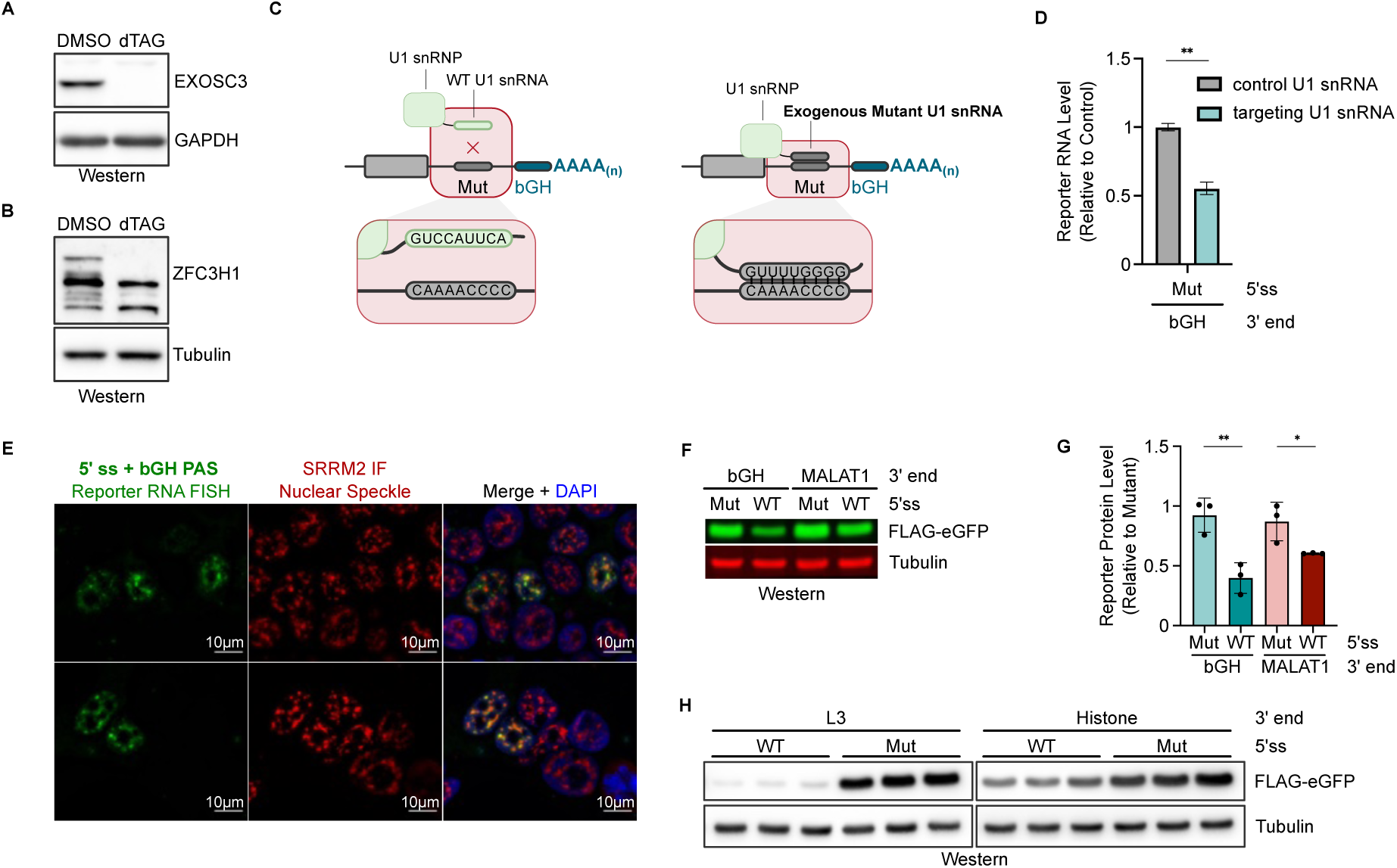
Additional Characterization of eGFP Reporters. **A**, Western blot analysis of EXOSC3 and GAPDH (loading control) levels in the EXOSC3 degron cell line that were mock-treated (DMSO) or dTAG-treated for 8 hours. **B**, Western blot analysis of ZFC3H1 and Tubulin (loading control) levels in the ZFC3H1 degron cell line that were mock-treated (DMSO) or dTAG-treated for 8 hours. **C**, Schematic of the U1 mutant snRNA expression approach used in **D**. Left: Wild-type (WT) U1 snRNP complexes will not recognize the 5′ ss Mut-*bGH* PAS reporter as the sequences are not complementary. Right: The exogenous mutant U1 snRNA will be loaded into U1 snRNP complexes, allowing U1 snRNP to recognize the *bGH* PAS-containing reporter with a Mut 5′ ss in its 3′ UTR. **D**, RT-qPCR of oligo d(T)-primed cDNA following overexpression of the reporter and the 5′ ss mutant-targeting U1 snRNA or a non-targeting U1 snRNA control. The reporter mRNA levels were first normalized to the expression of a co-transfected firefly luciferase control plasmid. The mRNA level of the reporter mRNA in the mutant-targeting U1 snRNA condition was then calculated relative to the non-targeting control U1 snRNA condition. Statistical analysis from *n* = 3 independent samples was calculated using an unpaired t-test. **E**, Representative images of reporter RNA FISH and SRRM2 immunofluorescence (IF) in wild-type HEK293T cells to monitor co-localization of the reporter RNA molecules and nuclear speckles. SRRM2 is used as a marker of nuclear speckles. The reporter RNA FISH signal is shown in green. SRRM2 IF signal is shown in red. Nuclei were stained using DAPI and are depicted in blue. **F**, Western blot analysis of FLAG-eGFP and Tubulin (loading control) levels after overexpressing the eGFP reporters with the *bGH* PAS or the *MALAT1* 3′ end in wild-type HEK293T cells for 48 hours. Proteins were detected using IR dye-conjugated secondary antibodies. **G**, Quantification of **F** using ImageQuant TL software. Each reporter protein level sample was first normalized to the loading control protein level. For each 3′ end pair, the relative protein levels were then calculated by dividing the reporter protein levels by one replicate of the 5′ ss mutant reporter. Statistical analysis of *n* = 3 independent samples was performed using unpaired t-tests. **H**, Western blot analysis of FLAG-eGFP and Tubulin (loading control) levels after overexpressing the eGFP reporters with the L3 PAS or the *H2AC18* (Histone) 3′ end in wild-type HEK293T cells for 48 hours. Three independent samples are shown for each reporter.

**Supplemental Figure 3.**
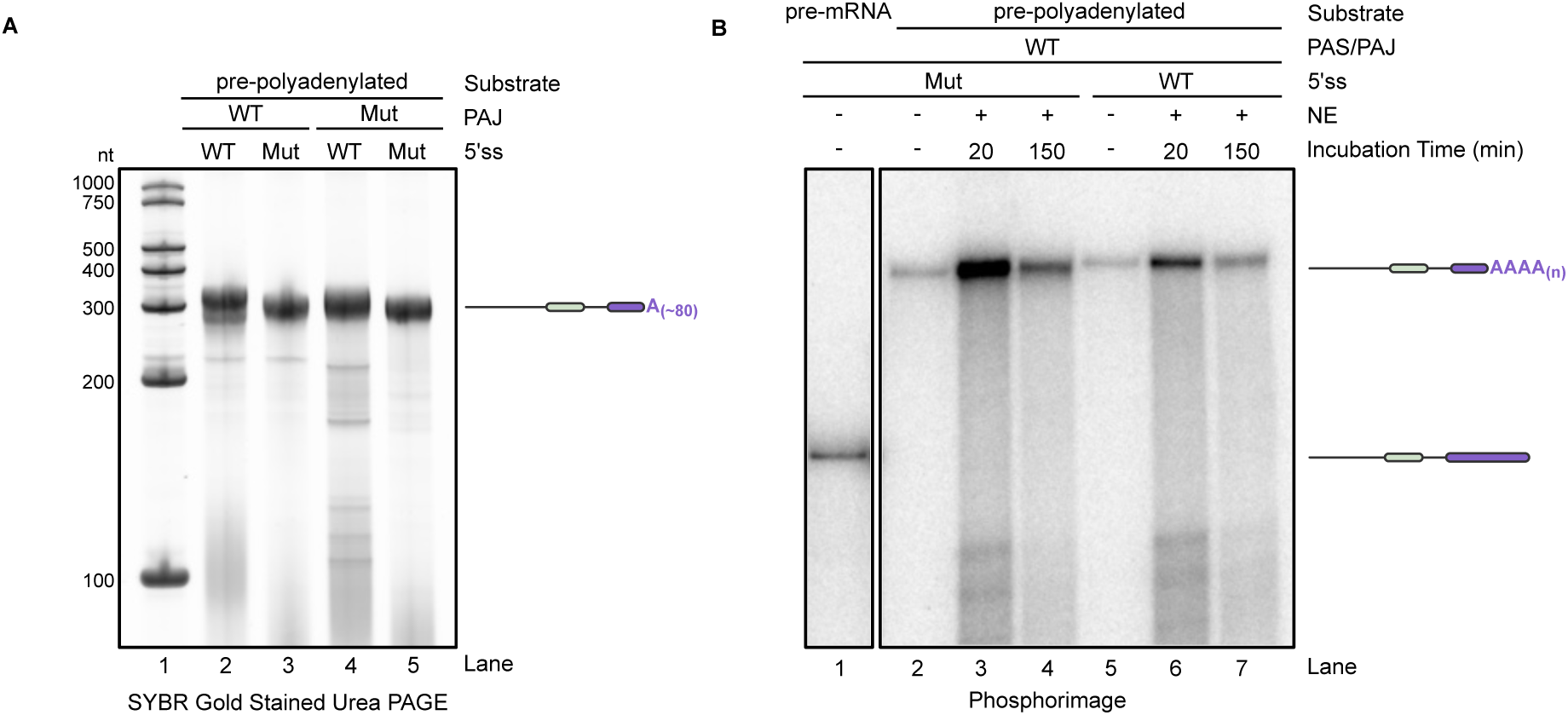
Characterization of pre-polyadenylated RNA substrates used in RNA pulldown assays. **A**, SYBR Gold stained urea-PAGE of pre-polyadenylated RNA substrates. **B**, Polyadenylation assay using radiolabeled pre-polyadenylated RNA substrates. The pre-mRNA lane (Lane 1) shown to provide a length comparison is from Lane 7 of Fig. 3, as all samples were resolved on the same gel.

**Supplemental Figure 4.**
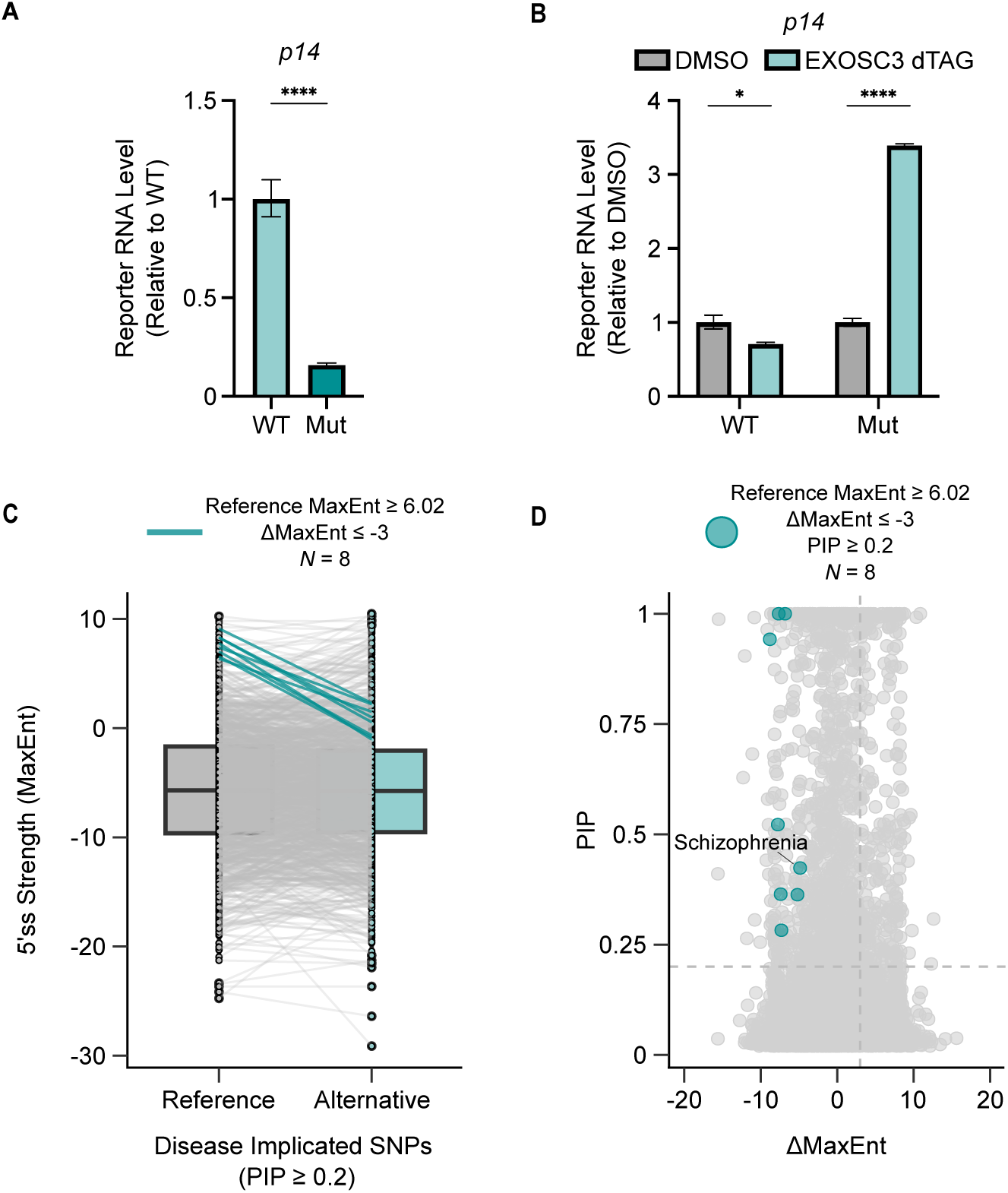
Disease-Associated SNPs and the Nuclear RNA Degradation Code. **A**, RT-qPCR analysis of oligo d(T)-primed cDNA following transfection of each reporter into HEK293T cells. Each reporter’s mRNA level was normalized to expression of a co-transfected firefly luciferase control. Expression of the *p14* Mut reporter was then calculated relative to the *p14* WT reporter. Statistical analysis from *n* = 3 independent samples was calculated using an unpaired t-test. **B**, RT-qPCR analysis of oligo d(T)-primed cDNA following overexpression of each reporter and dTAG-induced depletion of EXOSC3. The reporter mRNA levels were first normalized to the expression of a co-transfected firefly luciferase control plasmid. Then for each reporter, the mRNA level in the EXOSC3 dTAG condition was calculated relative to the control condition (DMSO). Statistical analysis from *n* = 3 independent samples was calculated using unpaired t-tests. **C**, Boxplot with paired points depicting the 5′ ss strength of all reference and alternative allele variants with a PIP score ≥ 0.2. Lines connect each reference and variant SNP MaxEnt score. Variants marked with a teal line have a reference MaxEnt score of at least 6.02 (1 IQR below the median 5′ ss strength across all 5′ splice sites) and the variant decreases the reference MaxEnt score by at least 3 MaxEnt units. **D**, Scatterplot depicting the change in MaxEnt score (ΔMaxEnt) and the PIP score for all variants. Variants marked with a teal circle have a PIP score of at least 0.2, have a reference MaxEnt score of at least 6.02 (1 IQR below the median 5′ ss strength across all 5′ splice sites), and the variant decreases the reference MaxEnt score by at least 3 MaxEnt units. The variant-associated disease is marked for one SNP.

**Supplemental Figure 5.**
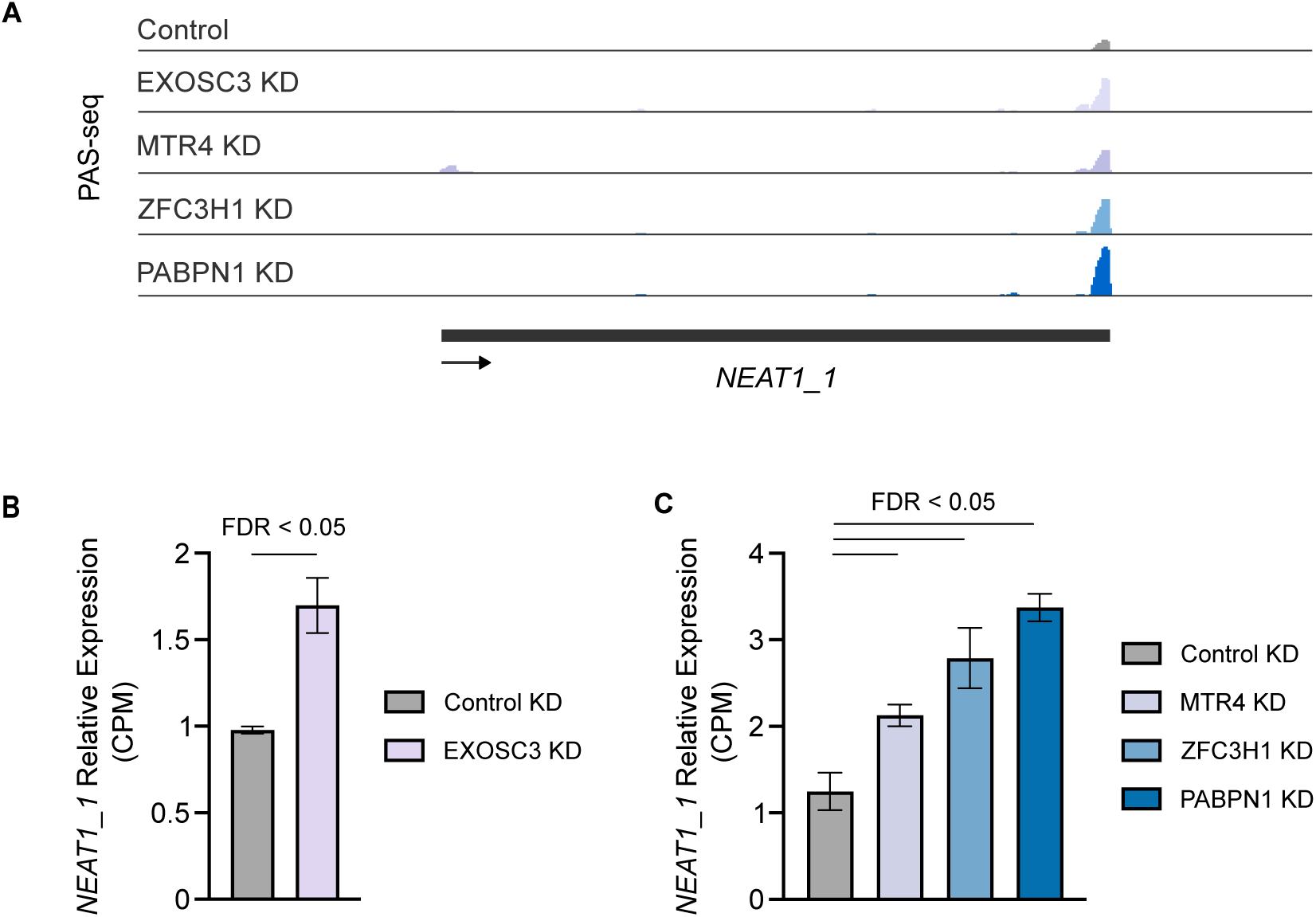
The short, polyadenylated isoform of *NEAT1* is upregulated following depletion of PAXT or the nuclear RNA exosome. **A**, PAS-seq data tracks for the gene *NEAT1_1* following depletion of EXOSC3, MTR4, ZFC3H1, or PABPN1 by RNAi in wild-type HEK293T cells. Each peak represents reads mapped to the 3′ end of mRNAs. **B** & **C**, Relative Expression (CPM: counts per million) of *NEAT1_1* as reported by differential isoform expression analysis of PAS-seq data from HEK293T cells depleted of EXOSC3 (**B**), MTR4, ZFC3H1, or PABPN1 (**C**) by RNAi. *NEAT1_1* exhibited at least a 1.5-fold increase in expression in all samples (log_2_FC > 0.585, FDR ≤ 0.05).

